# Transcriptional correlates of cocaine-associated learning in striatal ARC ensembles

**DOI:** 10.1101/2023.12.13.571585

**Authors:** Marine Salery, Arthur Godino, Yu Qing Xu, John F Fullard, Romain Durand-de Cuttoli, Alexa R LaBanca, Leanne M Holt, Scott J Russo, Panos Roussos, Eric J Nestler

**Affiliations:** Nash Family Department of Neuroscience & Friedman Brain Institute, Icahn School of Medicine at Mount Sinai; New York, NY 10029, USA; Department of Psychiatry & Friedman Brain Institute, Icahn School of Medicine at Mount Sinai; New York, NY 10029, USA; Department of Genetics and Genomic Sciences, Icahn Genomics Institute & Friedman Brain Institute, Icahn School of Medicine at Mount Sinai; New York, NY 10029, USA; Center for Disease Neurogenomics, Icahn School of Medicine at Mount Sinai; New York, NY 10029, USA; The Brain-Body Research Center of the Friedman Brain Institute, Icahn School of Medicine at Mount Sinai, New York, NY 10029; Mental Illness Research Education and Clinical Center (VISN 2 South), James J. Peters VA Medical Center, Bronx, 10468, NY, USA

## Abstract

Learned associations between the rewarding effects of drugs and the context in which they are experienced underlie context-induced relapse. Previous work demonstrates the importance of sparse neuronal populations – called neuronal ensembles – in associative learning and cocaine seeking, but it remains unknown whether the encoding vs. retrieval of cocaine-associated memories involves similar or distinct mechanisms of ensemble activation and reactivation in nucleus accumbens (NAc). We use ArcCreER^T2^ mice to establish that mostly distinct NAc ensembles are recruited by initial vs. repeated exposures to cocaine, which are then differentially reactivated and exert distinct effects during cocaine-related memory retrieval. Single-nuclei RNA-sequencing of these ensembles demonstrates predominant recruitment of D1 medium spiny neurons and identifies transcriptional properties that are selective to cocaine-recruited NAc neurons and could explain distinct excitability features. These findings fundamentally advance our understanding of how cocaine drives pathological memory formation during repeated exposures.

## Main Text

Cocaine use disorder remains a major public health challenge, with rates increasing in recent years in the absence of FDA-approved treatments. Cocaine produces its initial rewarding effects by increasing dopaminergic transmission in the nucleus accumbens (NAc), part of the ventral striatum. Adaptations in NAc neurons induced by repeated exposure to cocaine – at the transcriptional, cellular, and synaptic levels – have been related to specific behavioral abnormalities in rodent models that define an addiction syndrome (*1–5*). One striking finding, however, is that only sparse populations of NAc neurons respond to cocaine acutely or chronically based on the induction of immediate early genes (IEGs) (*6–12*). Identification of such cocaine-recruited “neuronal ensembles” could be analogous to ensembles identified in other brain regions where they are implicated in memory storage and retrieval (*13–19*) These ensembles of memory-associated neurons, referred to as engram cells, are thought to represent the memory trace associated with a given experience and are under intensive investigation (*20–22*).

IEGs are used to identify neuronal ensembles because their induction is coupled to synaptic activation (*23*) and because they serve as reliable proxies to characterize the spatiotemporal organization of such ensembles (*24–27*). Previous work using the IEG *Fos* has demonstrated the importance of NAc neuronal ensembles during relapse to cocaine seeking (*28–30*). However, it remains unclear whether the encoding and retrieval of cocaine-associated memories is supported by similar mechanisms of ensemble activation and reactivation that have been proposed in traditional learning paradigms. For example, it is unknown how individual cells that are activated by an initial exposure to cocaine respond over time after repeated cocaine exposures. It is also unknown whether the “acute” vs. “repeated” ensembles contribute to cocaine-associated memories in a similar or distinct manner. Further, while IEGs have been extensively used as markers of cellular activation to identify neuronal populations recruited by a specific stimulus, less attention has been paid to the lasting molecular consequences of their induction within ensemble cells, especially given that IEGs are postulated to be critical links between synaptic activity and downstream neuronal plasticity at both the transcriptional and synaptic levels.

Here, we address these crucial questions by leveraging one particular IEG, *Arc*. *Arc* encodes activity-regulated cytoskeleton-associated (ARC) protein, which has been shown to regulate several aspects of cocaine action within the NAc (*31–35*), making it an ideal IEG for identifying cocaine-elicited ensembles. Using transgenic ArcCreER^T2^ mice (*36*) crossed with various Cre-dependent reporter lines, we characterize how ARC ensembles shift over time during repeated cocaine exposure, with distinct functional effects on cocaine-linked memories. We also use single nucleus RNA-sequencing (snRNAseq) to not only determine the cellular constituents of acute vs. repeated ensembles but also the transcriptional adaptations that occur in ensemble cells over time.

## Cocaine-recruited ARC ensembles in NAc are reshaped by repeated cocaine exposure

As a first step in exploring the activation dynamics of NAc ARC ensembles, we measured endogenous ARC induction in response to acute or repeated cocaine exposure. Mice were habituated to a new context for 3 days, and then administered cocaine either acutely or repeatedly in this same context (Fig. 1, A). *Arc* mRNA induction, measured 1 h after the last injection, was significantly lower following repeated cocaine than acute cocaine (Fig. 1, B). Additionally, *Arc* mRNA levels positively correlated with locomotor responses after acute but not repeated treatment (Fig. 1, C). By comparison, *Fos* did not exhibit the same desensitization, and positive correlations between *Fos* mRNA expression and locomotor responses were found in both acute and repeated groups (fig. S1).

**Fig. 1:**
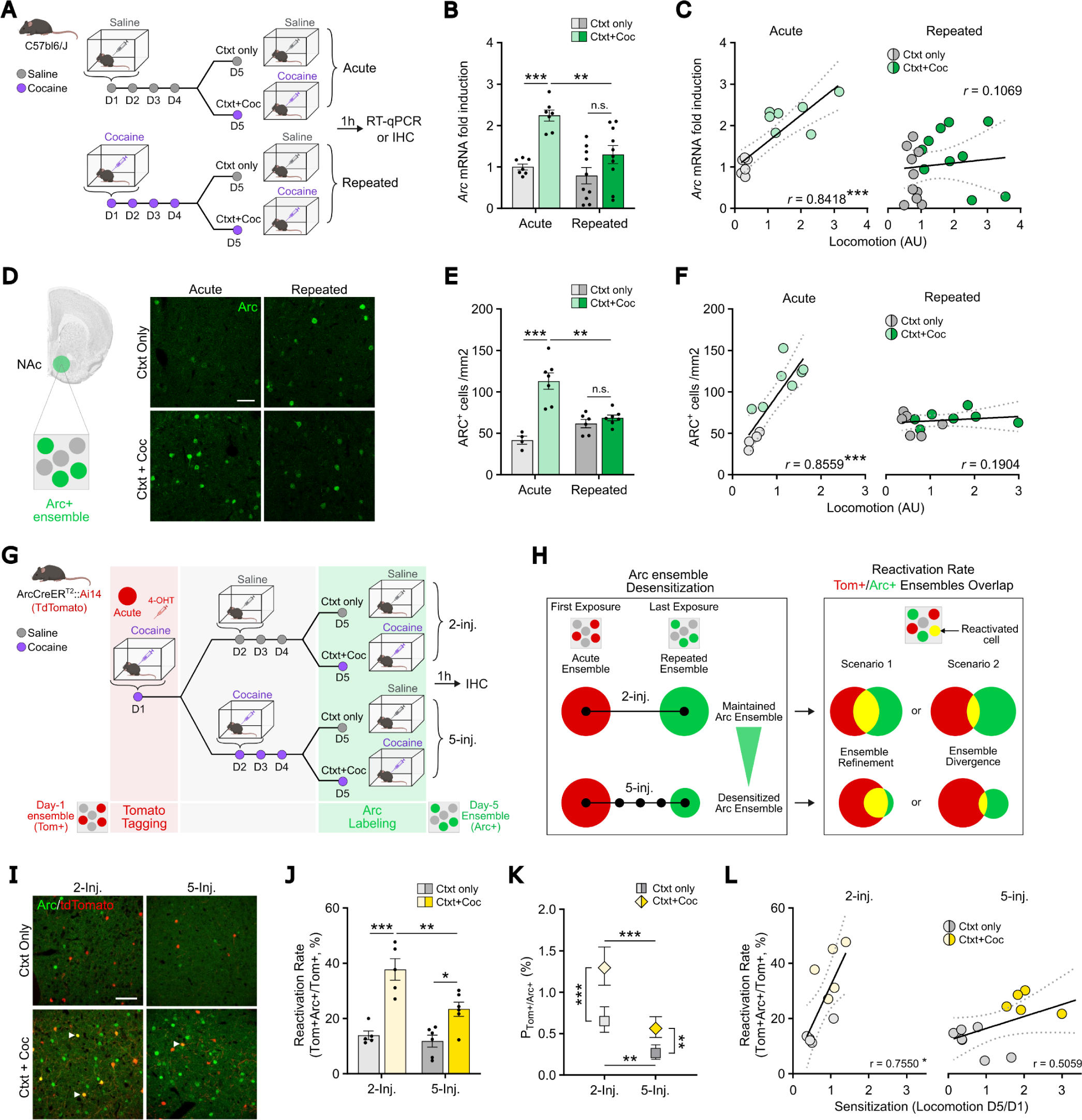
A smaller and mostly distinct ensemble is recruited after repeated cocaine-context associations. **(A)** Experimental design for acute vs. repeated experimenter-administered cocaine regimens in male C57BL/6J mice. Mice were injected i.p. with cocaine (20 mg/kg) or saline in an open-field arena different from their home-cage. On day 5, tissue was collected 1 h after the last injection. RT-qPCR: reverse transcription quantitative polymerase chain reaction, IHC, immunohistochemistry, Ctxt = context, Coc = cocaine. **(B)** RT-qPCR showed increased *Arc* mRNA levels in NAc upon acute, but not repeated, cocaine exposure. n = 7, acute; n=10, repeated. Two-way ANOVA: interaction regimen x drug, F_1,30_ = 3.897, *p* = 0.058; main effect of regimen, F_1,30_ = 9.577, \*\**p* = 0.0042; main effect of drug, F_1,30_ = 22.04, \*\*\**p* < 0.0001; followed by a Šidák correction for multiple comparisons. **(C)** *Arc* mRNA levels correlated positively with cocaine-induced locomotion in the acute (left), but not the repeated (right), group. Acute (left): n = 14, Pearson’s r = 0.8418, \*\*\**p* = 0.0002. Repeated (right): n = 20; Pearson’s r = 0.1069, *p* = 0.6536. **(D)** Representative confocal images of ARC+ cells (green, ARC+ ensemble) in NAc. Scale bar, 50 µM. **(E)** IHC quantification showed increased number of ARC+ cells in NAc upon acute but not repeated cocaine. n = 4, acute/ctxt only; n = 7, acute/ctxt+coc; n=6, repeated/ctxt only; n = 7, repeated/ctxt+coc. Two-way ANOVA: interaction regimen x drug, F_1,20_ = 21.04, \*\*\**p* = 0.0002; main effect of regimen, F_1,20_ = 3.050, *p* = 0.0961; main effect of drug, F_1,20_ = 31.07, \*\*\**p* < 0.0001; followed by Šidák post-hoc tests. **(F)** The number of ARC+ cells correlated positively with cocaine-induced locomotion in the acute (left), but not the repeated (right), group. Acute (left): n = 11; Pearson’s r = 0.8559, \*\*\**p* = 0.0008. Repeated (right): n = 13; Pearson’s r = 0.1904, *p* = 0.5333. **(G)** Experimental design for ensemble tagging in a 2-vs. 5-injection paradigm. On day 1 mice were injected i.p. with 4-hydroxytamoxifen (4-OHT, 10 mg/kg) concomitantly with cocaine (20 mg/kg) in an open-field arena different from their home cage. On days 2-5, animals received daily injections of saline or cocaine. On day 5, tissue was collected 1 h after the last injection. **(H)** Schematic representation of ARC ensemble desensitization after 5 cocaine exposures vs. 2 cocaine exposures (left) and of two possible scenarios for how the 2- and 5-injection (inj.) ensembles compare in cell identity (right). **(I)** Representative confocal images of Tom+ cells (red, Tom+ ensemble) activated on day 1 and ARC+ cells (green, ARC+ ensemble) activated on day 5 in the NAc. The arrows indicate representative reactivated neurons that were activated both on day 1 and day 5. **(J)** In both the 2-inj. and 5-inj. regimens, the reactivation ratio was significantly higher in the NAc of mice re-exposed to the context plus cocaine as compared to the context only, with a significantly lower reactivation level in the 5-inj. group. n = 5, 2-inj. groups; n = 6, 5-inj. groups. Two-way ANOVA: interaction regimen x drug, F_1,18_ = 5.296, \**p* = 0.0335; main effect of regimen, F_1,18_ = 9.591, \*\**p* = 0.0062; main effect of drug, F_1,18_ = 44.09, \*\*\**p* < 0.0001; followed by Šidák post-hoc tests. **(K)** Probabilities for a NAc cell to be activated on day 1 (Tom+, red) and on day 5 (ARC+, green) for each regimen (2- and 5-inj.) estimated with multinomial logistic regression to account for chance levels. The probability of reactivation was higher in mice re-exposed to the context plus cocaine as compared to the context only, with a significantly higher chance of reactivation in the 2-inj. group. Baseline-Category Logit Mixed Model (BCLogMM) equation for Tom+/ARC+ counts: interaction regimen x drug, z = 0.267, p = 0.79; main effect of regimen, z = -4.411, ***p < 0.0001; main effect of drug, z = 4.630, ***p < 0.0001; followed by Šidák post-hoc tests. **(L)** The reactivation ratio positively correlated with locomotor sensitization, expressed as a ratio of locomotion on day 5 over day 1, in the 2-inj. group (left) but not the 5-inj. group (right). 2-inj. (left): n = 10, acute; Pearson’s r = 0.7550, \**p* = 0.0116. 5-inj. (right): n = 12; Pearson’s r = 0.5059, *p* = 0.0933. Bar graphs are expressed as mean ± SEM with circles showing individual data. Correlation graphs show the regression line with a 95% confidence interval. Probability estimates are shown with their 95% confidence interval.

We next assessed the impact of this *Arc* desensitization on the size of ARC+ ensembles in the NAc in the same paradigm (Fig. 1, D). The number of cocaine-activated ARC+ cells was significantly lower in the repeated group, indicating the recruitment of a smaller ARC+ ensemble as compared to the one recruited by an acute cocaine injection (Fig. 1, E), consistent with the desensitization observed at the mRNA level (Fig. 1, B). Moreover, the size of the recruited ensemble was positively correlated with locomotor responses in the acute group but not in the repeated group (Fig. 1, F). We conclude that repeated sessions of cocaine exposure in a new context reshape ARC expression patterns and lead to a shift in ARC ensemble size that is behaviorally relevant. This shift resulted from repeated exposures to cocaine since a single re-exposure, 4 days after the initial one, did not alter ARC ensemble size or the correlation of ensemble size with cocaine-induced hyperlocomotion (fig. S2).

## ARC ensembles recruited at initial vs. later stages of cocaine-related learning are mostly distinct

Because ARC induction patterns shift during repeated cocaine exposure, with acute vs. repeated ARC ensembles exhibiting different sizes and differently correlating with cocaine-induced hyperlocomotion, we hypothesized that the two ensembles might present distinct features and encode distinct experiences associated with different (early vs. late) stages of context-reward associative learning. At the cellular level, this would be represented in the degree of overlap between these two ensembles: do they comprise the same or distinct NAc cells? To measure this experimentally, we capitalized on ArcCreER^T2^ mice (which induce Cre recombinase in an activity-dependent manner) crossed to Ai14 reporter mice (which express td-Tomato [Tom] in a Cre-dependent manner) to label cocaine-activated cells with the indelible expression of Tom upon tamoxifen injection (fig. S3, A). Importantly, Tom expression was absent without tamoxifen treatment, demonstrating negligible levels of leak expression (fig. S3, B, C, D). We leveraged this approach to quantify whether cells from the acute ensemble (recruited on the first day of training by the first injection of cocaine) are recruited again in the repeated ensemble (on the last day by a fifth injection of cocaine, Fig. 1, G). ArcCreER^T2^::Ai14 mice were treated with acute or repeated cocaine in a novel context as above, following a context-dependent locomotor sensitization protocol that heavily relies on NAc circuits and plasticity (*37*). Cells activated on day 1 (acute ensemble) were permanently tagged with Tom via the concomitant injection of tamoxifen and cocaine, whereas cells recruited on day 5 (repeated ensemble) were visualized via the induction of endogenous ARC expression 1 h after the last cocaine injection (Fig. 1, G).

We considered two possible outcomes (Fig. 1, H): 1) The acute ensemble could be largely recruited again by the last cocaine exposure (scenario 1, high overlap), which – because the repeated ensemble is smaller in size compared to the acute one – we refer to as a “refinement” of the ensemble. 2) Alternatively, the last cocaine exposure after repeated treatment might mostly not re-recruit the acute ensemble (scenario 2, low overlap), but rather incorporate new cells reflecting a “divergence” between the NAc ensembles encoding early vs. later stages of context-reward associative learning. The overlap between acute (red, Tom+) and repeated (green, ARC+) ensembles was visualized in NAc (Fig. 1, I) with reactivated cells (yellow, Tom+/ARC+) corresponding to cells recruited at both day 1 and day 5. The level of reactivation/overlap was measured as the number of double-positive (Tom+/ARC+) cells normalized to the total number of Tom+ cells.

We compared the extent of reactivation of the acute ensemble in response to a single cocaine dose four days later (2-injection condition, which does not induce ARC desensitization, fig. S2) vs. four subsequent consecutive days of cocaine injections (5-injection condition, which does induce ARC desensitization and a smaller ensemble size), with control mice receiving saline injections on day 5 to examine the effect of context re-exposure alone. We found significant reactivation of ensemble cells by cocaine, as compared to the context only, in both the 2- and 5-injection conditions (Fig 1, J). However, the cocaine-induced reactivation was significantly lower in the 5- vs. 2-injection groups (Fig. 1, J). Likewise, chance-adjusted probabilities of reactivation for individual Tom+/ARC+ cells belonging to both the initial and reactivated ensembles were significantly higher in both cocaine re-exposed groups as compared to context re-exposure only, but significantly decreased in the 5- vs. 2-injection condition (Fig. 1, K). This rules out the possibility that decreased reactivation in the repeated group results solely from decreased Tom+ ensemble size. These results indicate that the repeated ensemble recruited after five consecutive injections of cocaine overlaps less with the acute ensemble and incorporates more newly activated cells compared to the ensemble recruited by cocaine after just one prior injection, suggesting that ARC desensitization is likely accompanied by a shift in ARC-inducing cells towards a new, largely separate ensemble (scenario 2, Fig. 1, H). To evaluate the behavioral relevance of ensemble reactivation, we correlated the degree of reactivation to context-dependent locomotor sensitization in individual mice, and found a strongly positive correlation in the 2-injection group but not in the 5-injection group (Fig. 1, L).

Taken together, our data show that a subset (20-40%) of the ensemble recruited by the first exposure to cocaine becomes reactivated upon re-exposure to the drug 4 days later, and that the extent of this reactivation drops substantially with repeated drug exposures, when a greater fraction of new cells forms the ensemble. These findings indicate that mostly distinct ensembles are recruited at early vs. later stages of cocaine-context associative memory encoding and suggest that acute and repeated ensembles might support distinct phases of this learning.

## Acute vs. repeated ARC ensembles are differentially recruited during the retrieval of cocaine-context associative memories

We next studied whether, akin to other types of associative learning (*38, 39*), the retrieval of cocaine-context associative memories triggers the reactivation of ensembles previously activated during encoding, and if such retrieval-associated reactivation might differ between acute and repeated ensembles given their divergence during memory encoding (Fig. 1). To this end, we used a cocaine place preference (CPP) paradigm and evaluated whether NAc cells recruited in ArcCreER^T2^::Ai14 mice during cocaine-context conditioning are recruited again during subsequent memory recall. Encoding cells were permanently tagged (Tom+, red) via the concomitant injection of tamoxifen and cocaine during conditioning, whereas recall-activated cells were visualized using induction of endogenous ARC (ARC+, green) during the test session (Fig. 2, A).

**Fig. 2:**
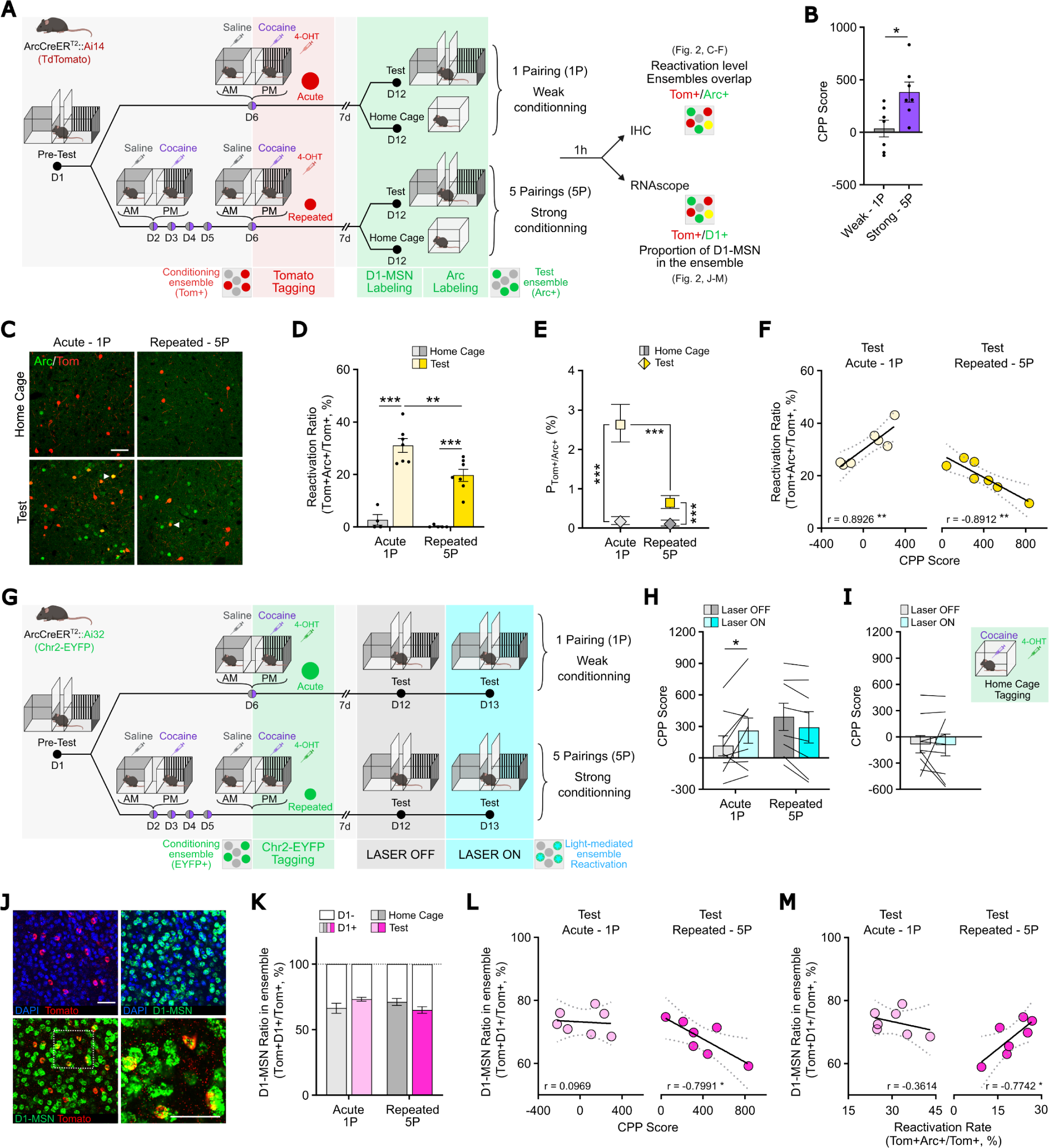
Acute and repeated ensembles differentially contribute to cocaine-context associative memories. **(A)** Experimental design for ensemble tagging in ArcCreER^T2^::Ai14 mice in a cocaine conditioned place preference (CPP) protocol. Mice were conditioned with cocaine (20 mg/kg i.p.) on one day or on five consecutive days and injected i.p. with 4-hydroxytamoxifen (4-OHT, 10 mg/kg) at the beginning of the last conditioning session. Mice were then tested 7 days later for their preference for the cocaine-paired chamber. NAc tissue was collected 1 h after the beginning of the test session. IHC, immunohistochemistry. **(B)** The repeated conditioning paradigm (5-pairing, 5P) induced a strong CPP score on the test day, whereas the acute conditioning (1-pairing, 1P) session failed to produce a place preference. n = 7, weak; n = 6, strong. Two tailed unpaired t test, \**p* = 0.0163. **(C)** Representative confocal images of Tom+ cells (red, Tom+ ensemble) activated during conditioning and ARC+ cells (green, ARC+ ensemble) activated during the test session. Arrows indicate representative reactivated neurons, i.e., cells activated during both conditioning and test. Scalebar, 50 µM. **(D)** In both the weak (1-pairing) and strong (5-pairing) conditioning paradigms, the reactivation ratio was significantly increased after the test session as compared to home cage controls, with a significantly lower reactivation in the 5-pairing group. n = 4, 1P/home cage; n = 5, 5P/home cage; n= 8, 1P/test; n = 10, 5P/test. Two-way ANOVA: interaction conditioning x test, F_1,19_ = 3.559, *p* = 0.0746; main effect of conditioning, F_1,19_ = 8.783, \*\**p* = 0.008; main effect of test, F_1,19_ = 104.9, \*\*\**p* < 0.0001; followed by Šidák post-hoc tests. **(D)** Probabilities for a NAc cell to be activated during conditioning (Tom+, red) and test (ARC+, green) for each conditioning paradigm (1- and 5-pairing) estimated with multinomial logistic regression to account for chance levels. The probability of reactivation was shown to be higher in mice subjected to the test as compared to the home cage controls, with a significantly higher chance of reactivation in the 1-pairing group. Baseline-Category Logit Mixed Model (BCLogMM), equation for Tom+/ARC+ counts: interaction conditioning x test, z = -2.093, *p = 0.0363; main effect of conditioning, z = -1.002, p = 0.3163; main effect of test, z = 8.984, ***p < 0.0001; followed by Šidák post-hoc tests. **(F)** The reactivation ratio was correlated positively with the CPP score in the 1-pairing group (left, positive correlation), but correlated negatively in the 5-pairing group (right, negative correlation). 1-pairing (left): n = 7; Pearson’s r = 0.8926, \*\**p* = 0.0068. 5-pairing (right): n = 7, Pearson’s r = - 0.8912, \*\**p* = 0.0071. **(G)** Experimental design for ensemble tagging and optogenetic-mediated reactivation in the CPP paradigm in ArcCreER^T2^::Ai32 mice. Mice were conditioned and given 4-OHT identically as shown in Panel A and 7 days later were tested for their place preference with the laser off and then on. **(H)** Light stimulation significantly increased the CPP score in the 1-pairing, but not the 5-pairing, group. n = 9, 1-pairing; n = 8, 5-pairing. Two-way ANOVA: interaction conditioning x light, F_1,15_ = 10.25, \*\**p* = 0.0059; main effect of conditioning, F_1,15_ = 0.8076, *p* = 0.3830; main effect of light, F_1,15_ = 0.3439, *p* = 0.5663. **(I)** ArcCreER^T2^::Ai32 mice were injected i.p. with 4-OHT plus cocaine in their home cage and the tagged ensemble was optogenetically-reactivated during the test session. Light stimulation of the home cage ensemble did not affect CPP scores. n = 8. Two tailed unpaired t test, *p* = 0.9576. **(J)** Representative confocal images of RNAScope staining showing the colocalization between Tom+ cells (red, Tom+ ensemble) activated during conditioning and D1+ MSNs. Scale bar, 50 µM. **(K)** RNAScope quantification showed a similar proportion of D1 MSN among the acute and repeated ensembles. n = 4, 1P/home cage; n = 5, 5P/home cage; n= 7, 1P/test; n = 10, 5P/test. Two-way ANOVA: interaction conditioning x test, F_1,22_ = 5.372, \**p* = 0.0302; main effect of conditioning, F_1,22_ = 03629, *p* = 0.5531; main effect of test, F_1,22_ = 0.0147, *p =* 0.9046; followed by Šidák post-hoc tests. **(L)** The ratio of D1 MSNs in the ensemble correlated negatively with the CPP score in the 5-pairing group (right), but not the 1-pairing group (left). 1-pairing (left): n = 7; Pearson’s r = - 0.0969, *p* = 0.8363. 5-pairing (right): n = 7, Pearson’s r = - 0.7991, * *p* = 0.0311. **(M)** The ratio of D1 MSNs in the ensemble showed significant positive correlation with ensemble reactivation in the 5-pairing group (right), but not the 1-pairing group (left). 1-pairing (left): n = 7; Pearson’s r = - 0.3614, *p* = 0.4257. 5-pairing (right): n = 7, Pearson’s r = 0.7742, * *p* = 0.0410. Bar graphs are expressed as means ± SEM with circles showing individual data. Correlation graphs show the regression line with a 95% confidence interval. Probability estimates are shown with their 95% confidence interval.

We first examined whether acute vs. repeated ensembles are more prone to reactivation during memory recall. We defined the level of recall-induced ensemble reactivation as the ratio of double positive Tom+/ARC+ cells to total Tom+ cells. The acute ensemble was captured upon an acute cocaine injection in a 1-pairing paradigm (1P, weak conditioning), while the repeated ensemble was captured on the last day of repeated cocaine injections in a 5-pairing paradigm (5P, strong conditioning, Fig. 2, A). As expected, repeated pairings induced a much stronger preference for the cocaine-paired side than an acute pairing, indicating more robust memory formation and a strong vs. weak conditioning with repeated vs. acute treatment, respectively (Fig. 2, B). Consistent with the relative desensitization of ARC induction after 5 daily doses of cocaine (Fig. 1, E), the repeated/strong-conditioning ensemble was smaller than the acute/weak-conditioning one (fig. S4), confirming the ability our of tagging approach to efficiently capture these two ensembles in a CPP paradigm.

Both acute and repeated ensembles exhibited significantly higher reactivation after the test session as compared to home cage controls, yet with the repeated ensemble being less reactivated than the acute one (Fig. 2, C, D). While the chance-adjusted probability for a cell to be reactivated during the test session was likewise increased for both ensembles, this probability was still significantly lower for cells belonging to the repeated/strong-conditioning ensemble (Fig. 2, E). We then asked whether the extent of an ensemble’s reactivation correlates with memory strength during the test session. We found opposite patterns: the acute ensemble’s reactivation was strongly positively correlated with CPP scores, whereas the repeated ensemble’s reactivation was strongly negatively correlated (Fig. 2, F). These data indicate that the recall of cocaine-context associative memories triggers the reactivation of a subset of cells that were previously recruited during the encoding of these memories, suggesting engram-like properties for NAc ARC ensembles in this context. Moreover, and somewhat counter-intuitively, the ensemble supporting a weak memory was more likely to be reactivated during recall, with the extent of its reactivation correlating positively with memory performance. Conversely, the comparatively lower reactivation of the strong memory-encoding ensemble was correlated negatively with memory strength. These results suggest opposite contributions of these two ensembles in the retrieval of cocaine-context associative memories.

To causally test this hypothesis, we used optogenetic-mediated neuronal activation to artificially promote reactivation of acute vs. repeated NAc ensembles during the retrieval of cocaine-context associative memories. Using the same cocaine CPP paradigm, acute and repeated ensembles were permanently tagged in ArcCreER^T2^::Ai32 mice, which express channelrhodopsin (ChR2-EYFP) in a Cre-dependent manner (fig. S5), thus allowing the permanent expression of ChR2 in conditioning-recruited cells and their subsequent light-induced reactivation during the test (Fig. 2, G). Experimental light-mediated reactivation of the acute, but not the repeated, ensemble elicited a higher CPP score during the test session as compared to a no-light, within-subject control condition (Fig. 2, H). As another control, reactivation of a cocaine ensemble captured in the home cage (where mice received equivalent exposures to cocaine without context conditioning) did not affect CPP performance (Fig. 2, I). These findings establish context-selective, yet differential roles for the acute and repeated ensembles and suggest that functionally distinct and mostly non-overlapping NAc cell populations cooperate to support both the encoding and the later retrieval of cocaine-context associative memories.

The NAc is composed of two mostly non-overlapping populations of medium spiny projection neurons (MSNs) that segregate between expression of D1 vs. D2 dopamine receptors and that have been shown to oppositely modulate CPP behavior (*40, 41*) We considered whether biased recruitment of D1 or D2 MSNs into the acute and repeated ensembles could explain the differences seen above. We thus quantified the proportion of tagged cells that express D1 receptors (D1+) vs. those that do not (D1-) using fluorescent *in situ* RNA hybridization (Fig. 2, J). First, acute and repeated ensembles exhibited a similar proportion (around 70%) of D1 MSNs (Fig. 2, K), suggesting that the portion of recruited D1 MSNs might not contribute to between-group functional differences. However, within-group ratios of D1 MSNs differentially correlated with CPP scores and with ensemble reactivation levels only for the repeated ensemble. A higher proportion of D1 MSNs in the repeated ensemble was associated with both a lower CPP score (Fig. 2, L) and higher reactivation level upon recall of cocaine-context memories (Fig. 2, M). Together, these data suggest that more subtle cell-specific features (within subpopulations of ARC+ D1 and D2 MSNs) might account for the differences observed between acute and repeated ensembles, rather than a distinct cell type composition.

## Ensembles recruited during acute vs. repeated encoding of cocaine-context associative memories have distinct plasticity-related transcriptional signatures

The differential contribution of the acute vs. repeated ensembles to memory retrieval could rely on divergent cellular properties, whether intrinsic or acquired through learning, coincident with or causal to *Arc* induction within ensemble cells. To more deeply phenotype the cells that constitute these ensembles, we combined snRNAseq with CPP testing to transcriptionally profile the individual NAc cells recruited in the Arc ensembles during the encoding and retrieval of cocaine-context associations by comparison to non-ensemble and to non-reactivated neighboring cells.

Similar to Tom tagging (Fig. 1, 2) and to ChR2 tagging (Fig. 2), acute and repeated conditioning ensembles were permanently tagged in ArcCreER^T2^::Sun1 mice, in which the permanent expression of a SUN1-GFP fusion protein allows for nuclei isolation via fluorescence-activated nuclei sorting (FANS, fig. S6). Tissue was collected 30 min after the beginning of the test session for the detection of recall-induced transcriptional programs in ensemble nuclei (Fig. 3, A). By combining each nucleus’s FANS status (GFP+/GFP-) and *Arc* mRNA expression (*Arc*+/*Arc*-), we segregated NAc nuclei and their transcriptional profiles according to their recruitment during, respectively, memory encoding (remote ARC induction, 7 days before retrieval) and memory recall (recent ARC induction, 30 min before sacrifice), or during both (reactivated, engram-like cells, GFP+/*Arc*+) (Fig. 3, A, B).

**Fig. 3:**
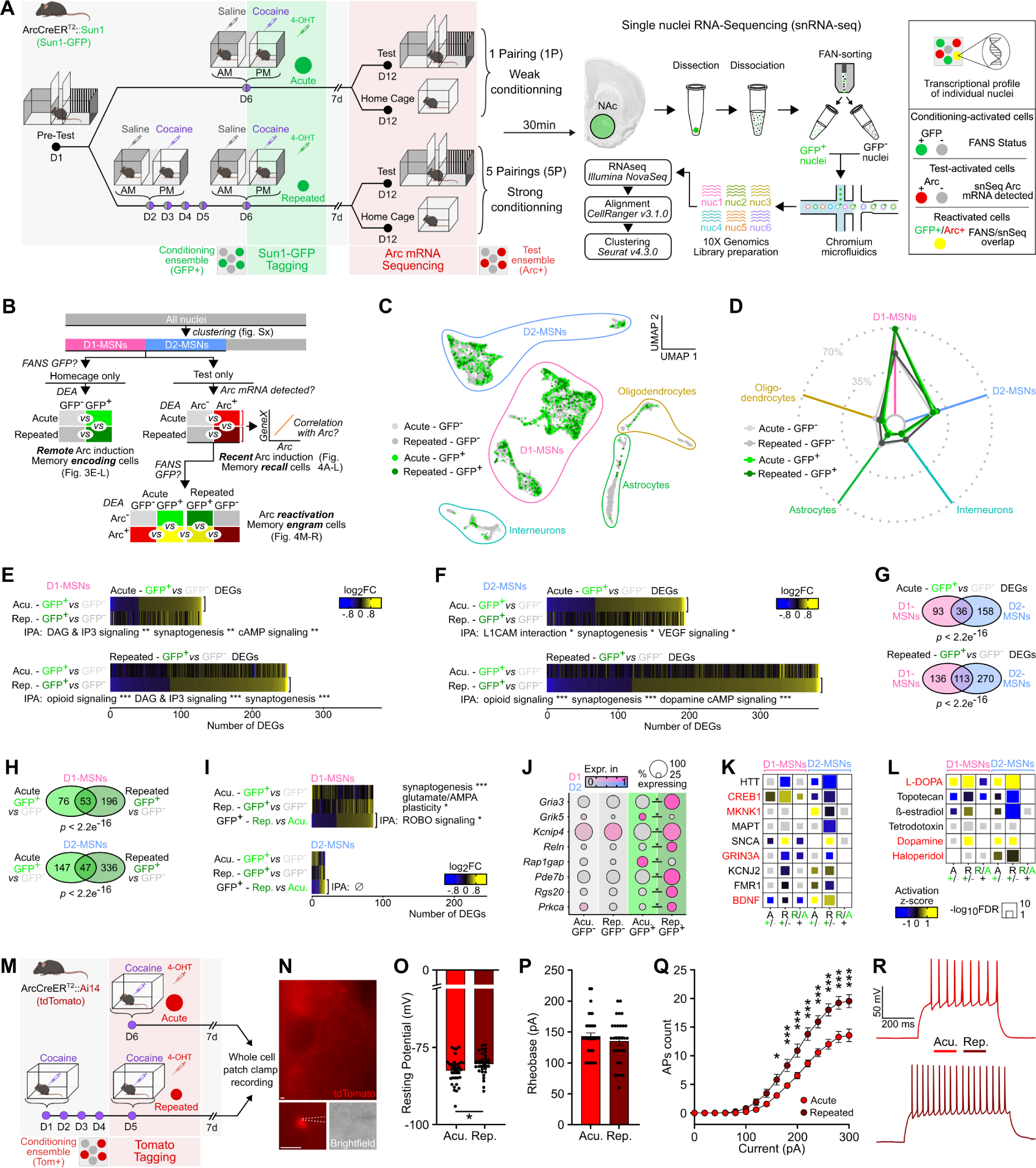
Transcriptional profiles of acute and repeated cocaine-context-associated ARC ensembles. **(A)** Experimental workflow for CPP ensemble tagging, FANS, and snRNAseq in ArcCreER^T2^::Sun1 mice. Mice were conditioned with cocaine (20 mg/kg i.p.) on one day or on five consecutive days and injected i.p. with 4-hydroxytamoxifen (4-OHT, 10 mg/kg) at the beginning of the last conditioning session. Mice were then tested 7 days later for their preference for the cocaine-paired chamber. NAc tissue was collected 30 min after the beginning of the test session and processed for snRNAseq. (**B**) snRNAseq analysis workflow for pairwise analyses of differentially expressed genes (DEGs). Home cage samples only were used for GFP+ vs GFP-comparisons to avoid any confounding effect of CPP testing, while CPP test only samples were used for ARC+ vs ARC-correlations. (**C**) UMAP reduction and cell type annotation of all collected nuclei (n = 11,539), colored according to treatment group and FANS status. (**D**) Proportion of nuclei from each treatment/FANS status combination in each cell type cluster, highlighting enrichment of GFP+ nuclei in D1 MSNs and depletion in glia. (**E**) Expression heatmaps of GFP+ vs GFP-DEGs in D1 MSNs in acute (top) and repeated (bottom) conditioning groups, along with corresponding 3 most significantly enriched IPA canonical pathways (** FDR < 0.01, *** FDR < 0.001). (**F**) Expression heatmaps of GFP+ vs GFP-DEGs in D2 MSNs in acute (top) and repeated (bottom) conditioning groups, along with corresponding 3 most significantly enriched IPA canonical pathways (* FDR < 0.05, *** FDR < 0.001). (**G**) Overlap of GFP+ vs. GFP-DEGs between D1 and D2 MSNs in either acute (top) or repeated (bottom) conditioning groups (Fisher’s exact test *p*-values). (**H**) Overlap of GFP+ vs GFP-DEGs between acute and repeated conditioning groups in D1 MSNs (top) and D2 MSNs (bottom) (Fisher’s exact test *p*-values). (**I**) Expression heatmaps of DEGs in GFP+ nuclei between repeated and acute groups in D1 MSNs (top) and D2 MSNs (bottom), along with corresponding 3 most significantly enriched IPA canonical pathways (* FDR < 0.05, *** FDR < 0.001). (**J**) Expression dot plots of selected D1 MSN DEGs involved in neuronal physiology and signaling (*FDR < 0.05). (**K**) Most significant IPA predicted upstream regulator proteins across comparisons in D1 and D2 MSNs. (**L**) Most significant IPA predicted upstream regulator chemical substances across comparisons in D1 and D2 MSNs. (**M**) Experimental design for ensemble tagging and whole-cell patch-clamp electrophysiological recordings in ArcCreER^T2^::Ai14 mice. Note that these cells (n = 34, acute; n = 30, repeated) are tagged with tdTomato (Tom+, red) but conceptually correspond to GFP+-tagged cells in the rest of the figure. (**N**) Representative wide-field (top) and close-up (bottom) views of patched MSNs. Scale bars 20 µm. (**O**) Resting membrane potential was less hyperpolarized in Tom+ neurons from the repeated ensemble. Two tailed unpaired t test, \**p* = 0.0473. (**P**) Rheobase, the minimal injected current required to trigger an action potential, was unaffected across groups. Two tailed unpaired t test, \**p* = 0.0473. (**Q**) The number of evoked action potentials (APs) in response to increasing depolarizing current steps was increased in Tom+ cells from the repeated ensemble compared to Tom+ cells from the acute ensemble, demonstrating enhanced excitability, Two-way ANOVA: interaction current x regimen, F_30,1120_ = 4.089, \*\*\**p* < 0.0001; main effect of current, F_15,1120_ = 124.1, \*\*\**p* < 0.0001; main effect of regimen, F_2,1120_ = 76.92, \*\*\**p* < 0.0001; followed by Šidák post-hoc tests. (**R**) Representative membrane responses from a Tom+ neuron in response to a 280 pA current injection from an acutely (top) or chronically (bottom) treated mouse. Bar and line graphs are expressed as means ± SEM.

From a merged dataset containing transcriptomes from all captured GFP+ and GFP-nuclei, unsupervised dimensionality identified 5 cell type clusters that were further annotated by comparison to publicly available snRNAseq databases (*42*) and that recapitulated well-characterized, canonical NAc cell types (fig. S7 and table S2). When quantifying the proportion of ensemble cells (GFP+) within each cell type, both acute and repeated ensembles showed a striking depletion in glial cells to the benefit of neuronal enrichment, with D1 MSNs accounting for the large majority (70%) of neurons in the ensemble cells (Fig. 3, C, D) – consistent with fluorescence *in situ* hybridization data (Fig. 2, J, K).

Next, we asked whether NAc cells allocated to the conditioning ensemble (GFP+) during memory encoding would exhibit persistent transcriptional features that distinguish them from cells that are not recruited in the ensemble (GFP-), and whether such features could further segregate acute vs. repeated ensembles at the molecular level. This first differential expression analysis was restricted to home-cage control samples to circumvent potential confounding effects of recall-induced transcription. Whether recruited in the acute or repeated ensembles, GFP+ cells separated from non-ensemble cells (GFP-) via the differential expression of numerous genes (table S3) associated with plasticity-related signaling pathways (e.g., dopamine cAMP signaling, synaptogenesis, Fig. 3 E, F). Importantly, GFP+ nuclei exhibited similar patterns of differentially expressed genes (DEGs) across ensembles (acute and repeated, Fig. 3, G) and cell types (D1 and D2 MSNs, Fig. 3, H), as shown by similar gene expression heatmap patterns and significantly overlapping gene lists. We then formally tested for statistically significant DEGs between acute and repeated GFP+ ensemble cells and found a higher number of DEGs in D1 than in D2 cells (Fig. 3, I). Furthermore, these repeated vs. acute DEGs were largely involved in synaptic function, signaling, and plasticity; a few select examples – glutamate receptors (*Gria3, Grik5*), ion channels (*Kcnip4*), and signal transducing proteins (*Reln, Rap1gap, Pde7b, Rgs20*) – are highlighted in Fig. 3, J. This confirmed that the acute and repeated ensembles, while sharing some common transcriptional properties (i.e., a shared transcriptional profile unique to all cocaine-activated cells and distinct from non-ensemble cells), also exhibit finer neuronal physiology-related differences that could underlie their distinct functional features. Finally, upstream regulator predictions confirmed that both acute and repeated ensembles engage transcriptional programs that are likely mediated by canonical striatal plasticity pathways (e.g., CREB, MAPK, BDNF signaling), and likely in a dopamine-dependent manner, in both D1 and D2 MSNs (Fig. 3, K, L).

We next assessed whether physiology-related transcriptional differences between acute and repeated ensemble cells have functional consequences. We thus examined the electrophysiological properties of these two ensembles in ArcCreERT2::Ai14 mice with *ex vivo* whole cell patch clamp recordings (Fig. 3, M, N). First, ARC+ cells recruited in the repeated ensemble exhibited a higher resting membrane potential (i.e., were less hyperpolarized at baseline) compared to acute ensemble cells (Fig. 3, O). Second, while rheobase was similar between the two ensembles (Fig. 3, P), cells from the repeated ensemble evoked significantly more action potentials with increasing steps of current injection (Fig. 3, Q, R), altogether indicating increased intrinsic excitability for repeated ensemble cells compared to acute ensemble cells.

Together, this initial snRNAseq analysis demonstrated that the activation of canonical striatal plasticity programs during cocaine-associated learning is largely restricted specifically to cells that previously induced ARC expression, underscoring at the single cell level that ARC induction represents not only a marker of recent activity, but also a predictor of later neuronal plasticity. This dataset also dissects the plasticity-related, activity- and dopamine-dependent transcriptional signature unique to cocaine-activated NAc cells. Beyond these shared features, we show as well that this molecular signature can discriminate between ensemble cells according to the stage of encoding at which they were recruited (early/acute ensemble vs. late/repeated ensemble), and that these transcriptional differences are associated with distinct electrophysiological properties in the two ensembles.

## Recent *Arc* induction denotes recall-evoked transcriptional regulation

As described above, the 1- vs. 5-pairing paradigms were associated with the expression of, respectively, weak or strong memory. We hypothesized that there are molecular correlates for such differences in memory strength, and asked whether the transcriptional programs engaged in recall-recruited cells would differ according to past cocaine history (1 vs. 5 pairing).

Cells recruited during the retrieval of cocaine-context associative memories were identified based on the detection of *Arc* mRNA within individual ensemble cells from CPP test samples only. Consistent with our previous findings (fig. S4), the number of *Arc+* expressing nuclei was increased by memory recall (test session, Fig. 4, A). With regard to cell types, a majority of recall-activated ensemble cells are D1 MSNs, although in slightly lower proportions than the encoding-recruited ensembles (Fig. 4, B, C). At the transcriptional level, cells recruited during recall (*Arc*+) were distinguished from non-recruited ones (*Arc*-) through the differential expression of numerous genes (table S4), largely involved in synaptic signaling and plasticity (e.g., synaptogenesis, dopamine cAMP signaling, MAPK cascade, Fig. 4, D, E). A significant proportion of these DEGs were regulated in both D1 and D2 MSNs (Fig. 4, F), illustrating that the transcriptional programs of recent *Arc* induction might exemplify a neuronal activation signature that goes beyond MSN subtype identity *per se*. Consistently, within each of these MSN populations, *Arc*+ cells also expressed higher levels of numerous other IEGs compared to *Arc*-cells, and furthermore *Arc* expression levels within individual nuclei were strongly correlated with those of the other IEGs (Fig. 4, G). Predicted upstream regulators of these gene sets implicated well-established activity-dependent transcriptional regulators, including CBP, as well as their likely dependence on dopamine signaling (Fig. 4, K, L). This finding both confirmed the ability of our snRNAseq approach to robustly identify nuclei with recent transcriptional activation, as well as the fact that those IEGs are induced together, in largely overlapping sets of NAc MSNs, upon CPP memory retrieval – which up to now has remained an outstanding question in the field. Despite significant similarities (Fig. 4, D, E, F), these recall-induced (*Arc*+) cells displayed significant gene expression differences depending on ensemble membership (acute, 1-pairing vs. repeated, 5-pairing groups), again implicating plasticity-related signaling pathways (e.g., CDK5 signaling, β-adrenergic signaling, Fig. 4, I). This last analysis points towards gene regulation mechanisms like “priming” and “desensitization” at the level of individual genes within individual cells: for instance, we highlighted genes that were only induced in activated cells with (e.g., *Kcnh5, Hcn1*) or without (e.g., *Npas2, Arhgef3, Tet2, Cacng3*) a history of repeated cocaine exposure (Fig. 4, J). Together, these data strongly support the idea that *Arc*-expressing ensemble cells support memory processes – including both memory encoding and retrieval – through activity-dependent plasticity- and memory-related transcriptional programs, and that these programs are modulated by past cocaine history (1 vs. 5 pairing).

**Fig. 4:**
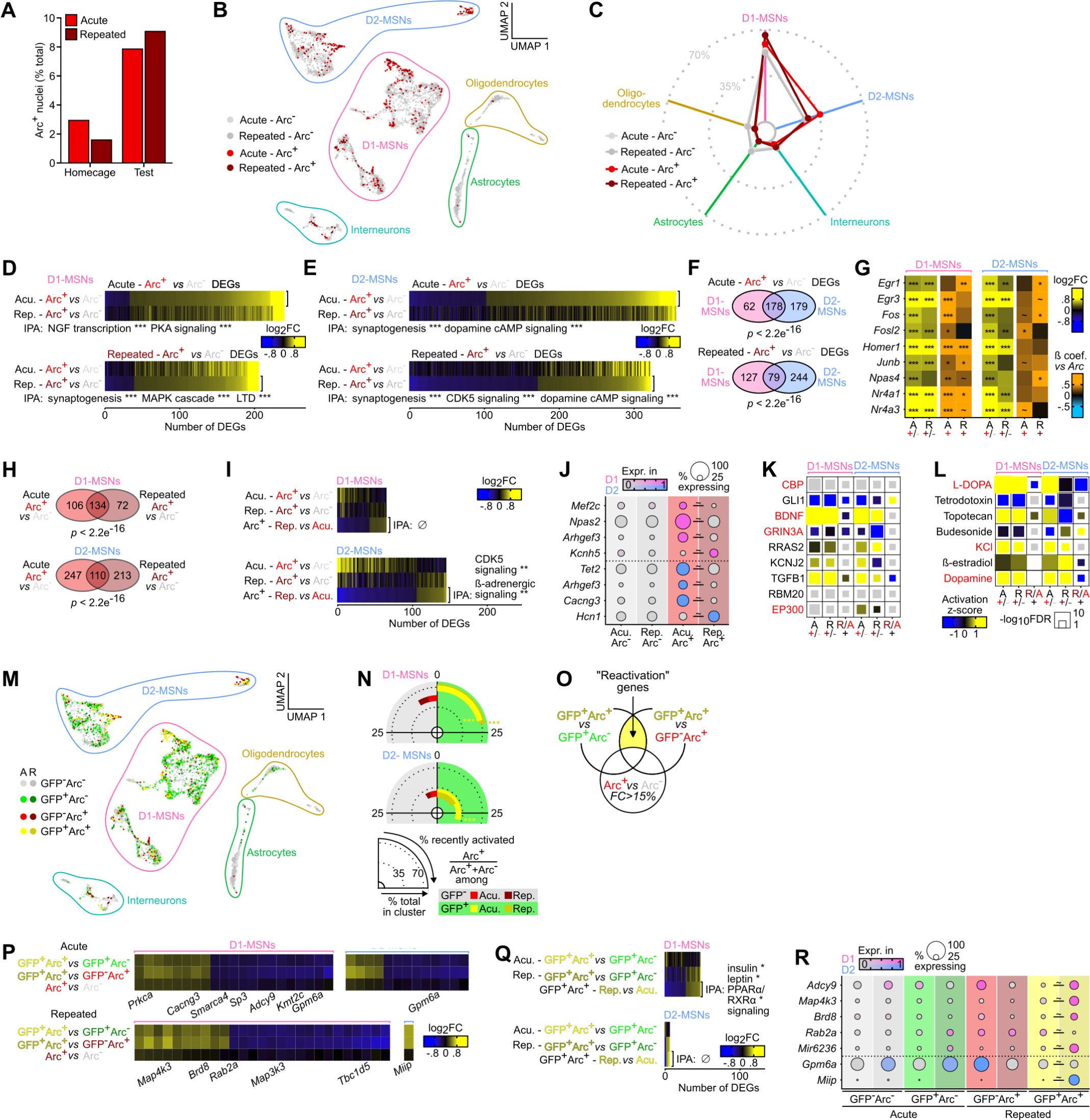
Transcriptional correlates of recent *Arc* induction and *Arc* reactivation in acute and repeated ensembles. **(A)** CPP testing increased the number of nuclei in which we detected *Arc* mRNA via snRNAseq, classified below as *Arc*+ nuclei, compared to home cage controls. (**B**) UMAP reduction and cell type annotation of nuclei from CPP test samples only (n = 5,448), colored according to treatment group and *Arc* expression status. (**C**) Proportion of nuclei from each treatment/*Arc* status combination in each cell type cluster, highlighting depletion in glia and enrichment in D1 MSNs. (**D**) Expression heatmaps of *Arc*+ vs *Arc*-DEGs in D1 MSNs in acute (top) and repeated (bottom) conditioning groups, along with corresponding 2 or 3 most significantly enriched IPA canonical pathways (*** FDR < 0.001). (**E**) Expression heatmaps of *Arc*+ vs *Arc*-DEGs in D2 MSNs in acute (top) and repeated (bottom) conditioning groups, along with corresponding 2 or 3 most significantly enriched IPA canonical pathways (*** FDR < 0.001). **(D)** Overlap of *Arc*+ vs *Arc*-DEGs between D1 and D2 MSNs in either acute (top) or repeated (bottom) conditioning groups (Fisher’s exact test *p*-values). **(G)** Expression and correlation heatmaps of selected IEGs, indicating concomitant broad IEG induction together with *Arc* in both D1 MSNs (left) and D2 MSNs (right). Correlation of IEG expression levels with *Arc* expression levels in individual *Arc*+ nuclei is shown. (**H**) Overlap of *Arc*+ vs *Arc*-DEGs between acute and repeated conditioning groups in either D1 MSNs (top) or D2 MSNs (bottom) (Fisher’s exact test *p*-values). (**I**) Expression heatmaps of DEGs in *Arc*+ nuclei between repeated and acute groups in D1 MSNs (top) and D2 MSNs (bottom), along with corresponding 2 most significantly enriched IPA canonical pathways (* FDR < 0.05, *** FDR < 0.001). (**J**) Expression dot plots of selected DEGs in D1 MSNs and D2 MSNs involved in neuronal physiology, signaling, and transcriptional regulation (∼FDR < 0.1). (**K**) Most significant IPA predicted upstream regulator proteins across comparisons in D1 and D2 MSNs. (**L**) Most significant IPA predicted upstream regulator chemical substances across comparisons in D1 and D2 MSNs. (**M**) UMAP reduction and cell type annotation of nuclei from CPP test samples only (n = 5,448), colored according to treatment group, GFP FANS status, and *Arc* expression status. (**N**) Proportion of nuclei from each treatment/GFP/*Arc* status combination in D1 MSNs (top) and D2 MSNs (bottom), highlighting preferential recent *Arc* induction (*Arc*+) in remote *Arc*-activated (GFP+) nuclei (D1, acute: χ^2^_(1)_ = 148.43, *p* < 2.2e^-16^; D1, repeated: χ^2^_(1)_ = 23.067, *p* = 1.56e^-6^; D2, acute: χ^2^_(1)_ = 102.77, *p* < 2.2e^-16^; D2, repeated: χ^2^_(1)_ = 2.57, *p* = 0.1083). (**O**) Schematic intersectional strategy to identify putative reactivation-related genes, i.e., genes that are DEGs between GFP+/*Arc*+ and GFP+/*Arc*-nuclei as well as between GFP+/*Arc*+ and GFP-/*Arc*+ nuclei but are not activity-dependent, i.e., are not changed overall in *Arc*+ vs. *Arc*-nuclei (for added stringency, all genes with an *Arc*+ vs. *Arc*-fold change >15% were removed, independent of FDR-based significance. (**P**) Expression heatmaps of putative “reactivation” genes in acute (top) and repeated (bottom) ensembles in both D1 MSNs (left) and D2 MSNs (right). Select genes involved in neuronal signaling and transcription are annotated. (**Q**) Expression heatmaps of DEGs in GFP+/*Arc*+ reactivated nuclei between repeated and acute groups in D1 MSNs (top) and D2 MSNs (bottom), along with corresponding most significantly enriched IPA canonical pathways (* FDR < 0.05). (**R**) Expression dot plots of selected D1 MSNs and D2 MSNs repeated vs. acute DEGs involved in neuronal physiology, signaling, and transcriptional regulation (∼ FDR < 0.1), that are also “reactivation” genes.

## ARC ensemble reactivation is transcriptionally mediated

One remaining question was to investigate whether these ensemble transcriptional features might play a part in governing reactivation at the single-cell level, especially given that we have demonstrated distinct reactivation properties for ensembles encoding distinct phases of learning (acute/1-pairing vs. repeated/5-pairing, Fig. 2). We hypothesized that NAc cells recruited in the acute vs. repeated ensembles during the encoding of cocaine-context associative memories acquire some divergent cellular properties that modulate their subsequent reactivation upon recall and consequently their contribution to memory retrieval. We thus examined whether cells reactivated during recall (*Arc*+/GFP+) exhibit different transcriptional responses depending on their former recruitment status, and whether such responses differ between acute vs. repeated ensemble cells.

Again, reactivated cells (*Arc*+/GFP+) are predominantly D1 MSNs (Fig. 4, M). Cells recruited during recall (*Arc*+) were significantly more likely to have been previously recruited in the encoding ensemble (GFP+), meaning that the recall ensemble preferentially recruited cells that were already recruited during encoding, except for D2 MSNs of the repeated group (Fig. 4, N). Such preferential re-recruitment of memory-encoding cells is consistent with earlier immunohistological observations above (Fig. 2, D-F) and with properties inferred for engram-like cells.

We then examined genes that were differentially regulated selectively in reactivated cells – by comparing cells activated during both encoding and recall (GFP+/*Arc*+) to cells only activated once during either encoding (GFP+/*Arc*-) or recall (GFP-/*Arc*+), and subtracting from the resulting DEG lists any genes that can be considered as activity-regulated (Fig. 4, O). Only a few genes survived this highly stringent selection process, for either D1 or D2 MSNs from either the acute or repeated ensemble (Fig. 4, P), yet with the potential to influence neuronal function at the levels of both synaptic signaling and transcriptional regulation. Finally, we formally examined gene expression differences between reactivated (GFP+/*Arc*+) cells from the repeated vs. acute ensembles (Fig. 4, Q and table S5) and found that “reactivation-predicting” genes appeared to also be DEGs in that last comparison (Fig. 4, R). This analysis suggests subtly distinct, transcriptionally-modulated reactivation mechanisms in the acute and repeated ensembles, which could account for their distinct reactivation probabilities (Fig. 2), and in turn could modulate their contribution at different stages of memory encoding.

## Discussion

The results of this study demonstrate that expression of a cocaine-context memory by re-exposure to the context triggers the reactivation of a subset of NAc neurons that had been previously activated during exposure to the drug itself and memory encoding, and that the extent of this reactivation correlates with memory strength. This ensemble of engram-like cells is likely to represent a cellular memory trace and a substrate for cocaine-associated memories. This is similar to what has been described for glutamatergic pyramidal neurons in hippocampus-related learning (*38, 39, 43*), only we are showing it here for the first time for GABAergic MSNs in NAc. It is striking to also note that the reactivation rates of these NAc cocaine engram cells appear similar if not higher than what tends to be observed for hippocampal engrams in Pavlovian learning paradigms (*38, 43*).

We go on to show that during a course of repeated cocaine exposures, the recruited NAc ensemble gets smaller but incorporates newly activated cells that were not recruited upon the first exposure. These findings establish that mostly distinct ensembles are recruited at early vs. later stages of cocaine-context learning. The differential contribution of these distinct neuronal populations to memory recall, which we demonstrate by use of optogenetic approaches, suggests that they could encode distinct aspects of these associative memories. The repeated ensemble – smaller, more excitable, and the reactivation of which during memory recall correlates negatively with memory strength – could exert a homeostatic effect on NAc circuits during later stages of learning. By contrast, our evidence shows that the acute ensemble drives memory formation and recall. Acute and repeated ensembles exist in brain coincidentally in time in response to a range of overlapping stimuli, and further work is needed to understand how their concomitant recruitment is integrated to determine behavioral responses.

Our snRNAseq dataset provides unprecedented insight into the transcriptional profiles of ensemble cells. We demonstrate for the first time that plasticity programs in NAc are largely restricted to cells that are functionally engaged in memory encoding and retrieval across cell types, and associate with *Arc* and other IEGs induction. By establishing the transcriptional correlates of ensemble allocation, we show that, beyond their activation at key phases of memory processes, ARC-defined ensemble cells also exhibit molecular features likely to support both their recruitment to the ensemble and their further contribution to memory storage through traditional plasticity mechanisms. Additionally, these data confirm that ARC expression efficiently stamps cell populations harboring transcriptional programs relevant to learning and memory. Future work is needed to establish which of these cellular adaptations are causal or consequential to ARC induction. Nevertheless, our data accentuate at single-cell resolution that ARC represents an effective marker of both recent neuronal activation but also of the induction of larger, longer-lasting, plasticity programs.

Finally, while we identify a shared molecular profile that distinguishes all ensemble-allocated cells (acute as well as repeated), we also provide evidence for more subtle transcriptional regulation that is specific to cells recruited at different stages of learning (i.e., acute vs. repeated ensembles). This finding indicates that transcriptional regulation tracks memory formation and storage at the cellular level. This work thereby provides more general insight into the mechanisms by which neuronal ensembles are shaped at the molecular level to control memory, and underscores the relevance of studying drug addiction as a form of pathological memory from both a fundamental and translational standpoint.

## Acknowledgments

The authors would like to thank Stephen Pirpinas, Katherine Beach, Catherine McManus, Kyra Schmidt, Nathalia Pulido, and Ezekiell Mouzon for transgenics breeding and genotyping, and Dr. Edgardo Aritzia from the Dean’s Flow Cytometry CoRE at the Icahn School of Medicine at Mount Sinai for his assistance with nuclei sorting. The authors would like to thank as well Dr. Nikos Tzavaras for his assistance with confocal image acquisitions that were performed at the Microscopy and Advanced Bioimaging CoRE at the Icahn School of Medicine at Mount Sinai.

## Author contributions

Conceptualization: MS, EJN

Methodology: MS, AG

Investigation: MS, AG, YX, RDC, AL, JFF, LH

Visualization: MS, AG

Funding acquisition: SJ, PR, EJN

Project administration: EJN

Supervision: EJN

Writing – original draft: MS, EJN.

Writing – review & editing: MS, AG, EJN

## Data and materials availability

All snRNAseq data reported in this study will be deposited in the Gene Expression Omnibus. *They will be made public upon publication - for review, please email the corresponding author for private access keys.* Custom R scripts and code utilized in this study, including for statistical analysis, are available upon request.

## Supplementary Materials

### Materials and Methods

#### Animals

C57BL/6J mice were purchased from The Jackson Laboratory at 7 weeks of age. ArcCreER^T2^ mice (IMSR_JAX:022357) were a generous gift from Christine Denny (*36*) at Columbia University and bred in-house on a C57BL/6J background. ArcCreER^T2^ mice were crossed in-house to the following transgenic reporter mouse lines obtained from the Jackson Laboratory: Ai14 (IMSR_JAX:007914), Ai32 (IMSR_JAX:012569), and Sun1 (IMSR_JAX:030952). All mice were maintained on a 12:12h dark/light cycle (08:00 lights off; 20:00 lights on) and were provided with food and water *ad libitum*. All behavioral testing was conducted in the dark phase at 8-18 weeks of age. All mice were maintained according to the National Institutes of Health guidelines for Association for Assessment and Accreditation of Laboratory Animal Care accredited facilities, and all experimental protocols were approved by the Institutional Animal Care and Use Committee at Mount Sinai.

#### Drug treatments

Cocaine HCl (from the National Institute on Drug Abuse) was diluted in 0.9% NaCl saline solution (ICU Medical) and injected intraperitoneally at 20 mg/kg. An aqueous formulation was used for 4-hydroxytamoxifen (4-OHT) delivery, as previously described (*44*). Briefly, 4-OHT (Sigma, H7904) was first dissolved in DMSO at 20 mg/mL, and then diluted to 1mg/mL in sterile saline containing 2% Tween-80. Mice were injected intraperitoneally with this solution at 10 mg/kg. For controls, the vehicle consisted of a saline solution only containing 2% Tween-80 and 5% DMSO, and was injected at a similar volume than 4-OHT (0.1 mL/10g body weight).

#### Activity-dependent tagging of ARC ensembles

Tagging of cocaine-activated cell populations was achieved with concomitant injection of cocaine and 4-OHT right before mice were placed into either the locomotor chamber (for locomotor sensitization) or the cocaine-paired chamber (for CPP). In both paradigms, mice were taken out of the chamber 1 h after the beginning of the behavioral task, put back into their home cages, and left undisturbed for 5 h to avoid any non-specific recombination and tagging.

#### Context-dependent locomotor sensitization

Locomotor activity was measured in a clear plexiglass open field arena (40 x 40 x 30 cm) as the number of laser beam-breaks collected over the duration of the test. Mice were placed in the open-field arena for 60 min after injection of a saline solution during three consecutive days for habituation before the actual experiment was performed. The protocol of context-dependent locomotor sensitization consisted of 5 daily sessions of 60 min in which locomotor activity was recorded for 60 min after an injection of saline or cocaine. Locomotor sensitization was calculated as the ratio of locomotion on day 5 over the locomotion on day 1.

#### Cocaine conditioned place preference

Unbiased cocaine conditioned place preference paradigm (CPP) was performed as previously described (*45*) using a three-chambered CPP set-up. Apparatus consisted of two chambers distinguished by distinct visual (gray vs. stripes walls) and tactile cues (small vs. large grid floors) separated by a small neutral area. Locomotion and time spent in each chamber was measured using an overhead video-tracking system (Ethovision XT 11) set to localize the mouse center point at each time of the trial. On the pre-conditioning phase (pretest day), mice were placed in the neutral area and allowed to freely explore the three chambers for 20 min. During the conditioning phase, drug-context learning was achieved by pairing an injection of saline with one chamber in the morning and a second injection of cocaine with the other chamber in the afternoon for either a single day (1-Pairing, weak conditioning group) or five consecutive days (5-Pairing, strong conditioning group). After injections, mice were confined to a given chamber for a period of 45 min. Groups and pairing sides were assigned after pretesting to balance any pre-existing chamber bias. On the post-conditioning phase, CPP testing was carried out with each mouse allowed again to freely explore all the chambers for 20 min. The CPP score was calculated as the difference in time spent on the cocaine-paired chamber vs. the saline-paired chamber during post-conditioning vs. pre-conditioning.

#### RNA extraction and quantitative real-time PCR

Mouse brains were collected after cervical dislocation and followed by rapid bilateral NAc punch dissections from 1 mm-thick coronal brain sections using a 14G needle and frozen on dry ice. RNA extraction was performed using the RNeasy Micro Kit (Qiagen) following manufacturer instructions. RNA 260/280 ratios of 2 were confirmed using spectroscopy, and reverse transcription was achieved using the iScript cDNA Synthesis 385 Kit (BioRad). Quantitative PCR using PowerUp SYBR Green (Applied Biosystems) was used to quantify cDNA using an Applied Biosystems QuantStudio 5 system. Each reaction was performed in triplicate and relative expression was calculated relative to the geometric average of 3 control genes (*Ppia, Tbp, Rpl38*) according to published methods (*45, 46*). Sequences of primers are available in table S1.

#### Immunohistochemistry (IHC)

At the indicated times after drug treatment or behavioral task (see figure legends), mice were rapidly anesthetized with an intraperitoneal injection of pentobarbital (50 mg/kg, Fatal Plus, Vortex Pharmaceutical) and transcardially perfused with a fixative solution containing 4% paraformaldehyde (PFA) (v/v) in 0.1 M phosphate buffer saline (PBS) delivered with a peristaltic pump at 6 mL/min for 5 min. Brains were removed from the skull and post-fixed for 24 h in a 4% PFA solution at 4°C. Sections of 30 µM thickness were cut in the frontal plane with a vibratome (Leica) and kept at -20°C in a cryoprotectant solution containing 30% ethylene glycol (v/v), 30% glycerol (v/v), and 0.1 M phosphate buffer (PB), Sections were then processed for IHC as previously described (*32*). Briefly, on day 1, sections were washed three times in 0.1 M phosphate buffer saline (PBS) and permeabilized for 15 min in a solution containing 0.2% Triton X-100 (Sigma Aldrich) in PBS. After three rinses in PBS, sections were incubated 1 h at RT in a blocking solution containing 10% Donkey Serum (v/v) in 0.1M PBS. Sections were then washed three times in PBS and incubated overnight at 4°C with the primary antibodies diluted in the blocking solution. The following primary antibodies were used either individually or in combination: ARC was detected using a rabbit polyclonal antibody raised against ARC (1/500, Synaptic System, 156-003), td-Tomato was detected using a goat polyclonal antibody raised against RFP (1/1000, Rockland, 200-101-379) and EYFP was detected using a polyclonal goat raised against GFP (1/1000, Aves Lab, GFP-1020). Following primary antibodies incubation, sections were rinsed three times in PBS and incubated 90 min at RT with adequate combination of the following secondary antibodies: donkey anti-rabbit Alexa-488-conjugated (1/500, Jackson Immunoresearch), donkey anti-goat Rhodamine Red (Jackson Immunoresearch) and donkey anti-chicken Alexa-488-conjugated (1/500, Jackson Immunoresearch). After three rinses in PBS, sections were incubated 10 min in DAPI for nuclei counterstaining before being rinsed three times in PBS, three times in PB and mounted in ProLong Diamond Antifade mounting medium (ThermoFisher Scientific).

#### RNA fluorescent in situ hybridization (FISH)

At the indicated times after drug treatment or behavioral task (see figure legends), mice were rapidly anesthetized with an intraperitoneal injection of pentobarbital (50 mg/kg, Fatal Plus, Vortex Pharmaceutical) and perfused transcardially using a fixative solution containing 4% paraformaldehyde (PFA) (v/v) in 0.1 M phosphate Buffer Saline (PBS) delivered with a peristaltic pump at 6 mL/min for 5 min. The brain was removed from the skull and tissue was post-fixed overnight in a 4% PFA solution and stored at 4°C. Sections of 30 µM thickness were cut in the frontal plane with a vibratome Leica, Nussloch, Germany) and kept at -20°C in a solution containing 30% ethylene glycol (v/v), 30% glycerol (v/v), and 0.1 M phosphate buffer. NAc slices were mounted on Superfrost Plus microscope slides (Fisher Scientific) and processed for RNA FISH using RNAscope Multiplex Fluorescent Reagent Kit v2 (ACD Bio) according to manufacturer instructions and as previously described (*45*) using mouse probes for *tdTomato* (tdTomato, #317041) and *Drd1a* (Mm-Drd1a-C2, #406491-C2) transcripts. Sections were counterstained with DAPI and mounted in ProLong Diamond Antifade mounting medium (ThermoFisher Scientific).

#### Images acquisition and quantification

For both IHC and FISH, images were acquired in a 16-bits range using a laser scanning confocal microscope (SP8, Leica) with a 40X oil immersion objective (Zoom 0.9, pixel size: x = 0.36 µm, y = 0.36 µm). Images were acquired in frame mode with a frame size of 1024 x 1024 pixels. Acquisition settings (laser intensities and gain) were kept identical across samples. Signal was quantified in NAc core and medial shell along the rostro-caudal axis for total of 12-15 different images (363.64 x 363.64 µm) per animal (i.e., 1 image per structure, per side, on three adjacent sections).

For IHC, the number of ARC-, tdTomato-, and/or EYFP-positive cells was manually counted in their respective channels using the “cell counter” plugin in Fiji. For each image, the total number of cells was quantified in the DAPI channel using a published custom pipeline (*45*). For single-channel quantifications, absolute numbers of ARC-, tdTomato-, and/or GFP-positive cells were normalized to the combined area imaged for each animal, and thus expressed as cells/mm^2^. For dual-channel quantifications (i.e., analyses of overlap or reactivation), the number of double positive cells was assessed by overlapping cell counter markers from each individual channel, and reactivation rates were calculated as: Reactivation 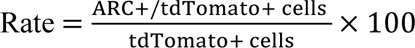. Further, we calculated reactivation levels over “chance” reactivation (i.e., the likelihood of double-positive cells given the respective number of single-positive cells for each channel among the total number of cells simply explained by random re-sampling), while accounting for the fact that these “chance” reactivation levels were unique to each animal (i.e., a nested design where individual cells are considered within individual animals). Animal-wise count data were treated as categorical with 4 outcomes (ARC-/tdTomato-, ARC-/tdTomato+, ARC+/tdTomato-, or ARC+/tdTomato+), and baseline-category logit models were fitted for these multinomial counts to estimate chance-adjusted log-odds for each outcome for each combination of group membership (2x2 design) using the *mclogit::mblogit* function and including a random effect for nesting. Estimated probabilities were then back-calculated from log-odds estimates. 95% confidence intervals for these estimated probabilities were computed by Monte-Carlo simulation, simulating 10,000 repeated draws from a multivariate-normal sampling distribution whose parameters are given by the logit model (logistic regression coefficients and variance-covariance matrix) with *MASS::mvrnorm*. Finally, group-wise probability estimates were compared with Sidak’s *post hoc* tests using *emmeans::emmeans*. These analyses were run in R v4.2.2. Custom R code and full model statistics are available upon request.

For FISH, the number of *tdTomato*- and/or *Drd1*-positive cells was manually counted in their respective channels using the “cell counter” plugin in Fiji (*47*). The number of double positive cells was assessed by overlapping cell counter markers from each individual channel, and D1 enrichment was calculated as: 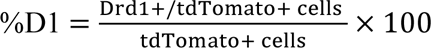.

#### Nuclei purification and fluorescence-activated nuclei sorting (FANS)

Mouse brains were collected after cervical dislocation and bilateral NAc dissection was rapidly performed from 1 mm-thick coronal brain sections using a 14G needle and tissue frozen on dry ice. Bilateral dissections from 16-20 mice were pooled together group-wise into one single sample. Nuclei suspension was obtained by homogenization of frozen pooled NAc samples in 4 mL of low-sucrose lysis buffer (0.32 M sucrose, 5 mM CaCl_2_, 3 mM Mg(Ace)_2_, 0.1 mM EDTA, 10 mM Tris-HCl) using a large clearance then a small clearance pestle of a glass Dounce tissue grinder (Kimble Kontes). The homogenates were filtered in an ultracentrifuge tubes (Beckman Coulter through a 40 μm cell strainer (Pluriselect) into ultracentrifuge tubes (Beckman Coulter), underlaid with 5 mL of high sucrose solution (1.8 M sucrose, 3 mM Mg(Ace)_2_, 1 mM DTT, 10 mM Tris-HCl). After centrifugation at 107,000 *g* for 1 h at 4°C in a SW41Ti Swinging-Bucket Rotor (Beckman Coulter, the supernatant was discarded, and nuclei pellets were resuspended in 800 µL of a solution containing 0.5% bovine serum albumin (BSA) in PBS and supplemented with DAPI at 1 µg/mL to allow for nuclei detection. Nuclei were sorted on a BD FACS Aria II three-laser device with a 100 μm nozzle and using BD FACSDiva Software v8.0.2. Gating strategies from a representative sort are visualized in fig. S7. Briefly, debris and doublets were excluded using FSC and SSC filters, nuclei were then selected as DAPI-positive (Violet1-A laser) events, and finally GFP-positive 610 nuclei (Blue1-A laser) were sorted directly into BSA-coated low-binding tubes. 15-18,000 nuclei were recovered for GFP+ samples and 100,000 for GFP-samples.

#### Single-nuclei RNA-sequencing (snRNAseq) and analysis

Following FANS, nuclei were quantified (Countess II, Life Technologies) and ±5,000 per GFP+ sample and ±20,000 per GFP-sample were loaded on a single 10X lane using Chromium Single Cell 3’ Library Construction Kit (10X Genomics). cDNA libraries were prepared according to the manufacturer’s protocol (10X Genomics). Libraries were sequenced at Azenta using the NovaSeq platform (Illumina) at a depth of ±350 million reads per sample. A Cell Ranger (v7.0.0) reference package was generated from the mm10 pre-mRNA mouse genome (GRCm38_v5) that ensured alignment to unspliced pre-mRNAs and mature RNAs. Cell Ranger filtered outputs were analyzed with Seurat v4.3.0 in R v4.2.2. Nuclei containing <900 reads, or <200 or >5000 features (i.e., genes for which at least one read was detected), or >1% of reads mapping to the mitochondrial genome were removed, leaving altogether 12,092 nuclei with a median 2,836 reads per nuclei for further analysis similarly to other previously published datasets (*48*). Nuclei from all samples then underwent integration using 3,000 features for *FindIntegrationAnchors*, clustering using 16 principal components and 20 nearest neighbors for *FindNeighbors* and a 0.1 resolution value for FindClusters following Seurat v4.3.0 vignette. These values were determined to recapitulate previously defined cell types. UMAP dimensionality reduction was finally run with *RunUMAP* on the *integrated_snn* graph calling the *r-reticulate* Python v3.6.10 install of *umap-learn v0.4.6* for visualization purposes. Libraries were normalized for size using *PrepSCTFindMarkers*, and marker genes for each cluster were computed with *FindAllMarkers* regressing out sample identity using logistic regression. Individual clusters were then further manually annotated by comparing enriched marker genes for each cluster (Supplemental fig. S7) with publicly available single-cell RNAseq databases of NAc tissue (*42*). Respectively 3 clusters of D1-MSNs and 2 clusters of D2-MSNs were manually combined together, and marker genes for each cell type cluster were computed with *FindAllMarkers* again (full marker gene lists and statistics are shown in table S2). One cluster of 553 glutamatergic nuclei was assumed to result from dissection contamination from the piriform cortex just anterior to NAc (which itself does not contain glutamatergic neurons), and corresponding nuclei were thus removed from further analysis. Libraries from the remaining 11,539 nuclei with a median 2,782 reads per nuclei were re-normalized for library size using *PrepSCTFindMarkers* again.

Cluster-specific pairwise differential expression analysis (a schematic of all comparisons is shown in Fig. 3) was performed using *FindMarkers* on SCT normalized counts. Fold change was computed for all genes, but statistical testing using logistic regression for differentially expressed genes (DEGs) was further restricted to genes detected in >30% of nuclei in at least one of the two groups in the corresponding pairwise comparison, and with >15% expression change between groups. *p*-values were adjusted for false discovery rate (FDR) at a 0.1 significance level (full DEG lists and statistics are provided in tables S3, S4, S5). A given nucleus was considered *Arc*+ if it contained at least one read mapping to *Arc* transcript. For comparisons between *Arc*+ and *Arc*-nuclei (Fig. 4), the FANS status of individual nuclei (GFP+/-) was regressed out as a latent variable to diminish its influence on differential expression. Within *Arc*+ nuclei, correlations between individual *Arc* counts and those of other transcripts was computed using linear regression (*stats::lm*), regressing out FANS status again, and signed ß coefficient estimates were extracted to determine the strength of the association between *Arc* and other genes’ expression. Preferential reactivation (Fig. 4) was assessed via χ^2^ testing of group-wise FANS-GFP status x *Arc* status contingency tables using *stats::chisq.test*. Enrichment analyses of gene expression patterns (significant DEG lists) was then performed using the Ingenuity Pathway Analysis (IPA, Qiagen) knowledgebase and classification system, extracting both enriched canonical signaling pathways and predicted upstream regulators significant by Fisher’s enrichment test statistical testing and FDR correction. Custom R scripts are available upon request.

#### Optogenetics

Male and female ArcCreER^T2^ mice were anesthetized with an intraperitoneal bolus of ketamine (100 mg/kg) and xylazine (10 mg/kg), then head-fixed in a stereotaxic apparatus (Kopf Instruments). Then, 200-µm-wide optical fibers (Doric, MFC_200/240-0.22_4.5mm_MF1.25_FLT) were bilaterally implanted above NAc at AP +1.5 mm, ML +1.5 mm, DV –4.0 mm, at a 10° angle. Optical fibers were secured in place using dental cement (3M) without the use of screws to the skull and covered with a layer of black dental cement (C&B Metabond). Mice were allowed >3 weeks to recover before experimentation. For optogenetic stimulation, mice were connected to a dual optical patch-cord (Doric) connected to a 473 nm blue laser (OEM Laser System). Stimulation was executed as trains of 10 box pulses (15 ms, 8-10 mW) emitted at 20 Hz every 5 s during the entire 20 min CPP test session. Individual animals were tested both with and without light illumination in a within-subject design.

#### *Ex vivo* whole-cell patch-clamp recordings

Mice were anesthetized using isoflurane. Brains were rapidly extracted, and coronal sections (250 mm) were prepared using a Compresstome (Precisionary Instruments Inc.) in cold (0-4°C) sucrose-based artificial cerebrospinal fluid (SB-aCSF) containing 87 mM NaCl, 2.5 mM KCl, 1.25 mM NaH_2_PO_4_, 4 mM MgCl_2_, 23 mM NaHCO_3_, 75 mM sucrose, 25 mM glucose. After recovery for 60 min at 32°C in oxygenated (95% CO_2_/ 5% O_2_) aCSF containing 130 mM NaCl, 2.5 mM KCl, 1.2 mM NaH_2_PO_4_, 2.4 mM CaCl_2_, 1.2 mM MgCl_2_, 23 mM NaHCO_3_, 11 mM glucose, slices were kept in the same medium at room temperature for the rest of the day and individually transferred to a recording chamber continuously perfused at 2-3 mL/min with oxygenated aCSF. Patch pipettes (4-6 MΩ) were pulled from thin wall borosilicate glass using a micropipette puller (Sutter Instruments) and filled with a K-gluconate (KGlu)-based intra-pipette solution containing 116 mM KGlu, 20 mM HEPES, 0.5 mM EGTA, 6 mM KCl, 2 mM NaCl, 4 mM ATP, 0.3 mM GTP (pH 7.2). Cells were visualized using an upright microscope with an IR-DIC lens and illuminated with a white light source (Olympus for Scientifica), and fluorescence visualized through a mCherry bandpass filter upon LED illumination through the objective (p3000ULTRA, CoolLed) using MicroManager v2.0 (https://micromanager.org/). All recordings were made on anterior NAc core MSNs. Excitability was measured in current-clamp mode by injecting incremental steps of current (0–300 pA, +20 pA at each step). For recording of spontaneous Excitatory Post-Synaptic Currents (sEPSCs), neurons were recorded in voltage-clamp mode at -70 mV and sEPSCs detected with a 8 pA threshold. Whole-cell recordings were performed using a patch-clamp amplifier (Axoclamp 200B, Molecular Devices) connected to a Digidata 1550 LowNoise acquisition system (Molecular Devices). Signals were low pass filtered (Bessel, 2 kHz) and collected at 10 kHz using Axon pCLAMP 11 Software Suite (Molecular Devices). Electrophysiological recordings were extracted using Clampfit (Molecular Devices). All groups were counterbalanced by days of recording and all recordings were performed blind to experimental conditions.

#### Statistics

No statistical power estimation analyses were used to predetermine sample sizes, which instead were chosen to match numerous previous publications (*32, 40, 45, 49*). Unless specified otherwise above (multinomial logistic regression and snRNAseq), all statistics were performed in GraphPad Prism v9. In summary, pairwise comparisons were performed with Welch’s *t*-tests, correlations using Pearson’s *r* and multifactorial designs were analyzed with ANOVAs with repeated measures when appropriate. Pairwise post-hoc testing was adjusted using Sidak’s correction. Bar and line graphs represent mean ± sem. Correlation graphs represent regression line with its 95% confidence interval. Significance was set at *p* < 0.05.

**Fig. S1:**
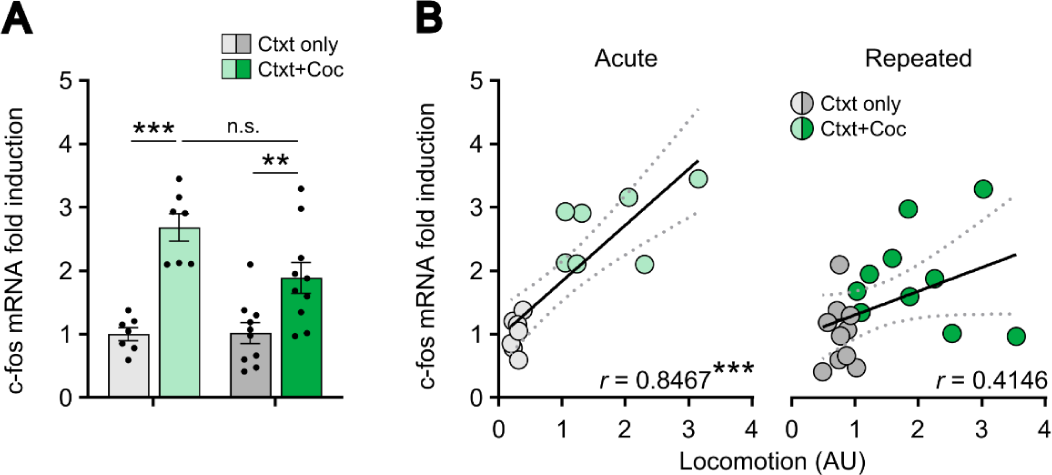
*Fos* expression is not desensitized after repeated cocaine exposure. **(A)** *Fos* mRNA levels in NAc are increased by both acute and repeated cocaine as compared to saline, without significant desensitization of *Fos* induction in the repeated group. Results are expressed as fold increase normalized to the Acute/Ctxt only group. n = 7, acute groups; n=10, repeated groups. Two-way ANOVA: Interaction regimen x drug, F_1,30_ = 4.032; *p* = 0.0537; main effect of regimen F_1,30_ = 3.715, *p* = 0.0634; main effect of drug F_1,30_ = 40.27, \*\*\**p* < 0.0001; followed by a Šidák correction for multiple comparisons. **(B)** *Fos* mRNA levels correlated positively with cocaine-induced locomotion in the acute (left), but not the repeated (right), group. Acute (left): n = 7, Pearson’s r = 0.8467, \*\*\**p* = 0.0001. Repeated (right): n = 10; Pearson’s r = 0.4146, \*\**p* = 0.0691. Bar graphs are expressed as means ± SEM with circles showing individual data. Correlation graphs show the regression line with a 95% confidence interval.

**fig. S2:**
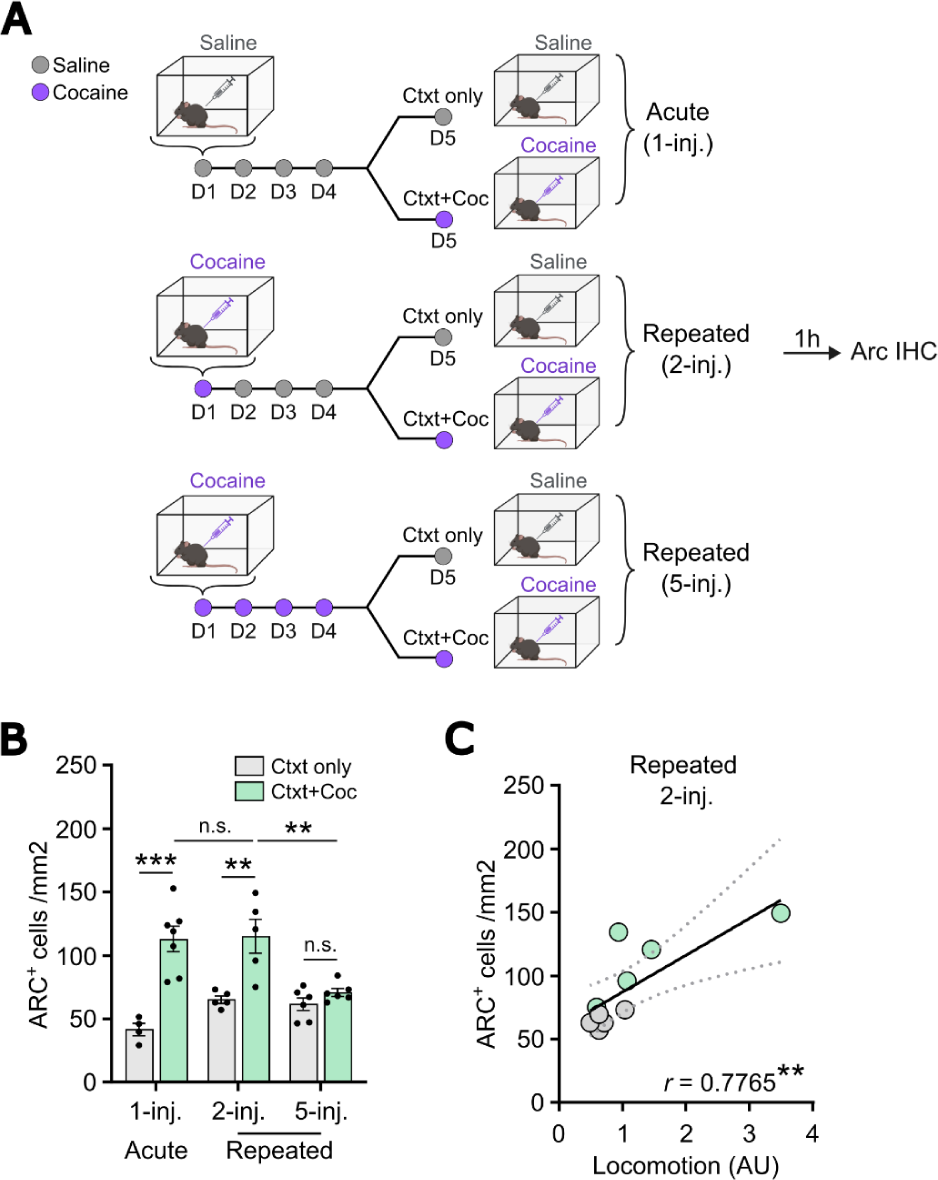
A single re-exposure to cocaine is not associated with a decrease in ARC ensemble size. **(A)** Experimental design for experimenter-administered cocaine regimens in C57BL/6J mice. Acute and repeated regimens are compared with a single cocaine re-exposure where animals are injected twice with cocaine 4 days apart. Mice were injected i.p. with cocaine (20 mg/kg) or saline in an open-field arena different from their home cage. On day 5, tissue was collected 1 h after the last injection. IHC, immunohistochemistry, Ctxt = context, Coc = cocaine, 1-inj = 1 injection, 2 inj. = 2 injections, 5-inj. = 5 injections. **(B)** The repeated 2-inj. group showed an increased number of ARC+ cells (green, ARC+ ensemble) in NAc after cocaine as compared to saline, to a similar extent as the acute 1-inj. group. Cocaine-mediated ARC induction in the 2-inj. group was significantly higher than in the repeated 5-inj. group. n = 4, acute/ctxt only; n = 7, acute/ctxt+coc; n=5, 2-inj./ctxt only and 2-inj./ctxt+coc; n=6, 5-inj./ctxt only; n = 7, 5-inj./ctxt+coc. Two-way ANOVA: interaction regimen x drug, F_2,27_ = 8.266; \*\**p* = 0.0016; main effect of regimen F_2,27_ = 4.642, \**p* = 0.0185; main effect of drug F_1,27_ = 44.88, \*\*\**p* < 0.0001; followed by Šidák post-hoc tests. **(C)** The number of ARC+ cells correlated positively with cocaine-induced locomotion in the 2-inj. group. Pearson’s r = 0.7765, \*\**p* = 0.0083. Bar graphs are expressed as means ± SEM with circles showing individual data. Correlation graphs show the regression line with a 95% confidence interval.

**fig. S3:**
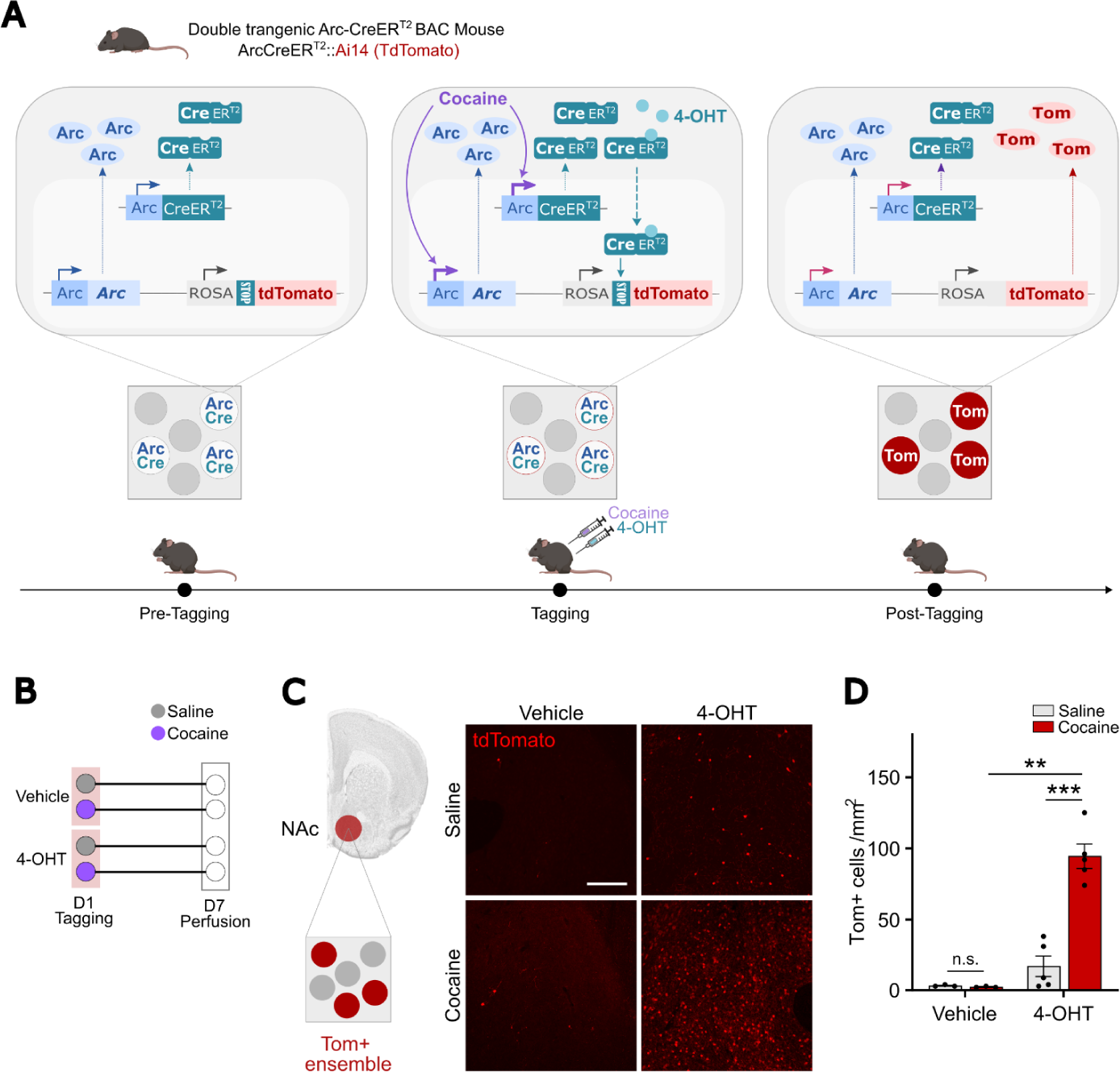
Permanent tagging of ARC+ ensembles in ArcCreER^T2^::Ai14 mice. (**A**) Schematic representation of the strategy applied to tagging cocaine-recruited ARC+ ensembles in NAc. ArcCreER^T2^ mice were crossed with the Ai14 reporter line to induce stable tdTomato expression in *Arc*-expressing cells. Ensemble tagging is achieved via the concomitant injection of 4-OHT and cocaine. Cocaine triggers induction of both endogenous ARC and the CreER^T2^ transgene. Upon tamoxifen binding, CreER^T2^ enters the nucleus where it removes the floxed-STOP cassette and enables tdTomato expression. The persistent expression of tdTomato in cocaine-activated cells allows for their visualization at any future time (e.g., 7 days). Tom, tdTomato; 4-OHT, 4-Hydroxytamoxifen. (**B**) On day 1 mice were injected i.p. with cocaine (20 mg/kg) or saline along with 4-hydroxytamoxifen (4-OHT, 10 mg/kg) or vehicle in their home cage. Tissue was collected 7 days later to visualize the ensembles previously recruited by cocaine. (**C**) Representative confocal images of Tom+ cells (red, Tom+ ensemble) in NAc. Scalebar, 100 µM. (**D**) As compared to saline, cocaine increased the number of Tom+ cells in the 4-OHT-treated group, but not in the vehicle-treated one. n = 3, Vehicle; n = 5, 4-OHT. Two-way ANOVA: Interaction treatment x pre-treatment, F_1,12_ = 27.48, ***p = 0.0002; main effect of treatment, F_1,12_ = 26.09; ***p = 0.003; main effect of pre-treatment, F_1,12_ = 49.98, ***p < 0.0001. Bar graphs are expressed as means ± SEM with circles showing individual data.

**fig. S4:**
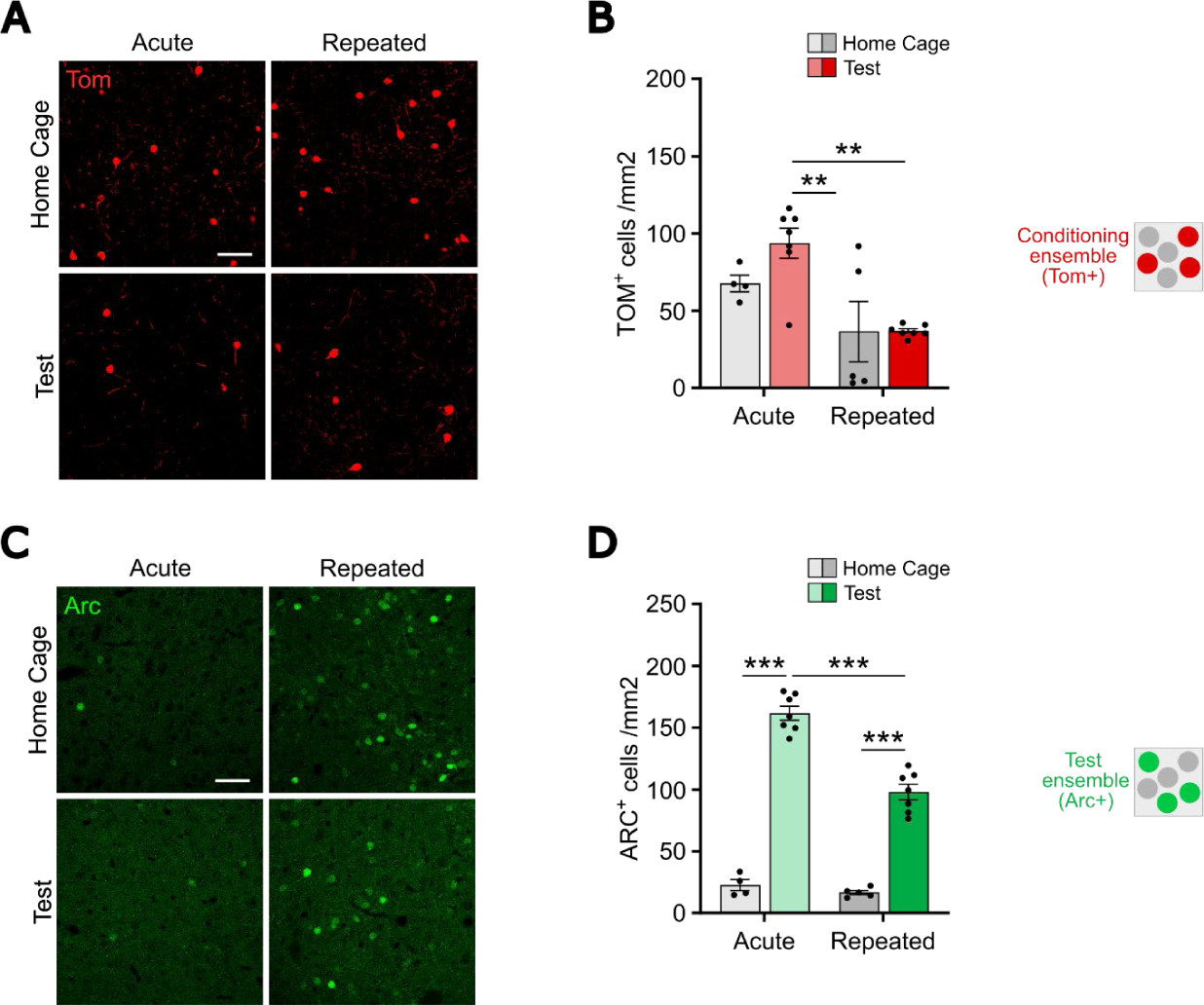
Recruitment of NAc ensembles during weak vs. strong conditioning and expression of cocaine-associated memories. **(A)** Representative confocal images of the tdTomato (Tom+) cells (red, conditioning ensemble) that are recruited in NAc during the conditioned phase of the cocaine conditioned place preference (CPP) paradigm. **(B)** IHC quantification showed a significantly higher number of cells recruited in the 1-pairing vs. 5-pairing groups without any effect of the test as compared to the home cage. n = 4, acute/home cage; n = 7, acute/test; n = 5, repeated/home cage; n = 7, repeated/test. Two-way ANOVA: interaction conditioning x test, F_1,19_ = 1.440, *p* = 0.2448; main effect of conditioning, F_1,19_ = 16.73, \*\*\**p* = 0.0006; main effect of test, F_1,19_ = 1.522, *p* = 0.2323; followed by a Šidák correction for multiple comparisons. **(C)** Representative confocal images of the ARC+ cells (green, test ensemble) recruited in NAc during the test session in the CPP protocol. **(D)** The number of ARC+ cells increased after the test session as compared to the home cage condition in both the 1-pairing and 5-pairing groups, with a significantly lower induction in the 5-pairing group. n = n = 4, acute/home cage; n = 7, acute/test; n = 5, repeated/home cage; n = 7, repeated/test. Two-way ANOVA: interaction conditioning x test, F_1,19_ = 26.29, \*\*\**p* < 0.0001; main effect of conditioning, F_1,19_ = 3851, \*\*\**p* < 0.0001; main effect of test, F_1,19_ = 3845, \*\*\**p* < 0.0001; followed by a Šidák correction for multiple comparisons.

**fig. S5:**
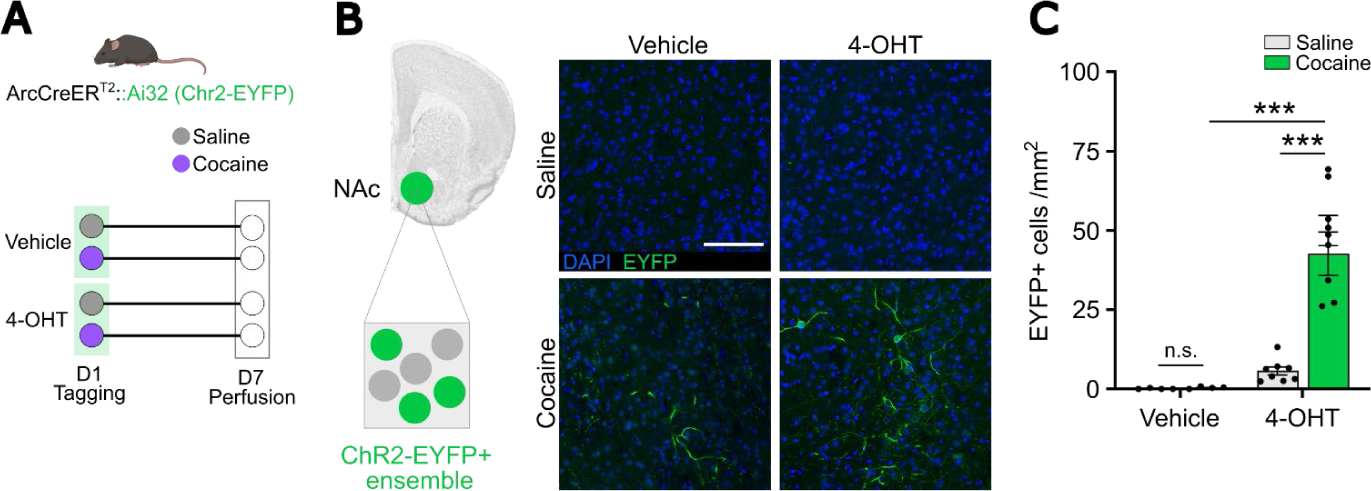
Permanent tagging of ARC+ ensembles in ArcCreER^T2^::Ai32 mice. (**A**) On day 1, mice were injected i.p. with cocaine (20 mg/kg) or saline along with 4-hydroxytamoxifen (4-OHT, 10 mg/kg) or vehicle in their home cage. Tissue was collected 7 days later to visualize the ensembles previously recruited by cocaine. (**C**) Representative confocal images of EYFP+ cells (green, ChR2-EYFP+ ensemble) in NAc. Scalebar, 100 µM. (**D**) As compared to saline, cocaine increased the number of EYFP+ cells in the 4-OHT-treated group, but not in the vehicle-treated group. n = 4, Vehicle; n = 8, 4-OHT/Saline, n = 6, 4-OHT/Cocaine. Two-way ANOVA: Interaction treatment x pre-treatment, F_1,18_ = 20.34, ***p = 0.0003; main effect of treatment, F_1,18_ = 21.04; ***p = 0.00; main effect of pre-treatment, F_1,18_ = 34.50 ***p < 0.0001. Bar graphs are expressed as means ± SEM with circles showing individual data.

**fig. S6:**
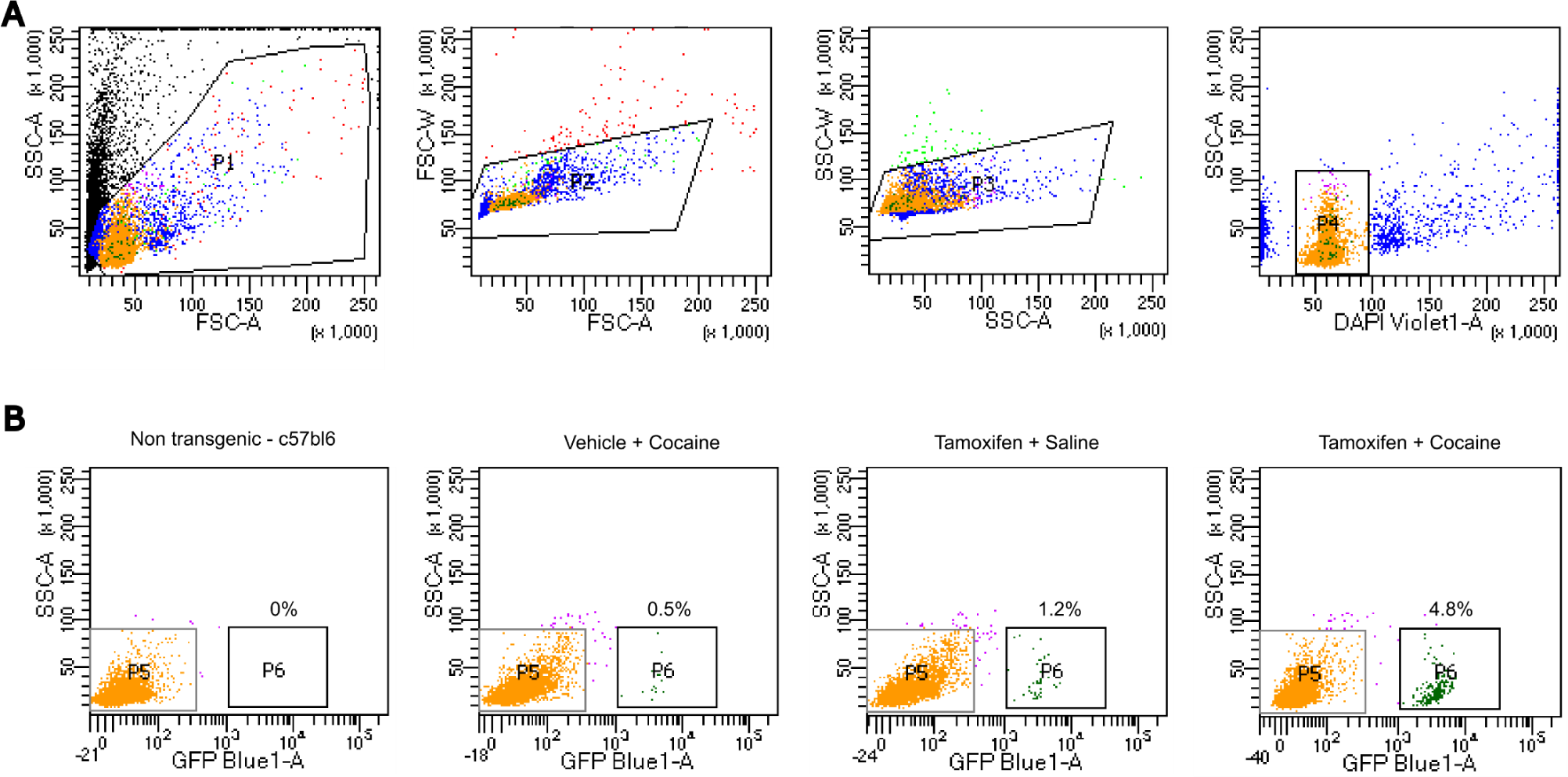
Fluorescence-Activated Nuclei Sorting (FANS) of striatal ARC neuronal ensembles. (**A**) Representative FANS gating strategy from a ArcCreERT2::Sun1 NAc sample. (**B**) Visualization of FANS-isolated non-ensemble (GFP-) and ensemble (GFP+) nuclei and percent of GFP+ nuclei for non-transgenic sample or ArcCreERT2::Sun1 treated with a combination of tamoxifen (or vehicle) along with cocaine (or saline). Tamoxifen (or vehicle as a control) was injected concomitantly with cocaine (or saline as a control) in the home cage and NAc collected 7 days later for nuclei isolation and FANS-based ensemble isolation.

**fig. S7:**
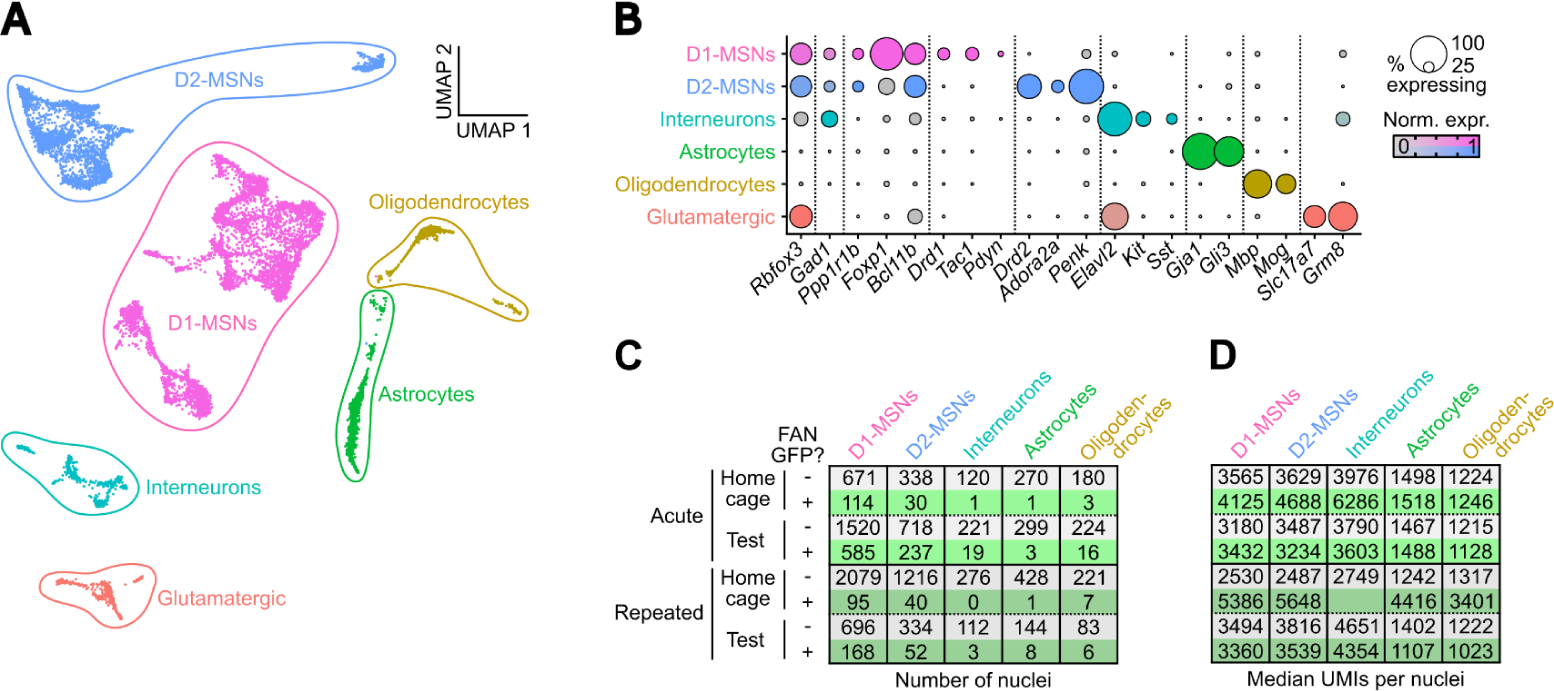
Cell type annotation of snRNAseq clusters. (**A**) UMAP reduction and cell-type annotation of all collected nuclei (n = 11,539), segregated into phenotypically defined clusters and colored according to their cluster of origin. (**B**) Expression of published marker genes *(42)* for striatal cell types across clusters. Full list of cluster marker genes is given in table S2. (**C**) Number of nuclei from each group and FANS status for each cell type. (**D**) Median number of unique molecular identifiers (UMIs) detected in nuclei from each group and FANS status for each cell type.

### Supplementary Tables

**Table S1:**
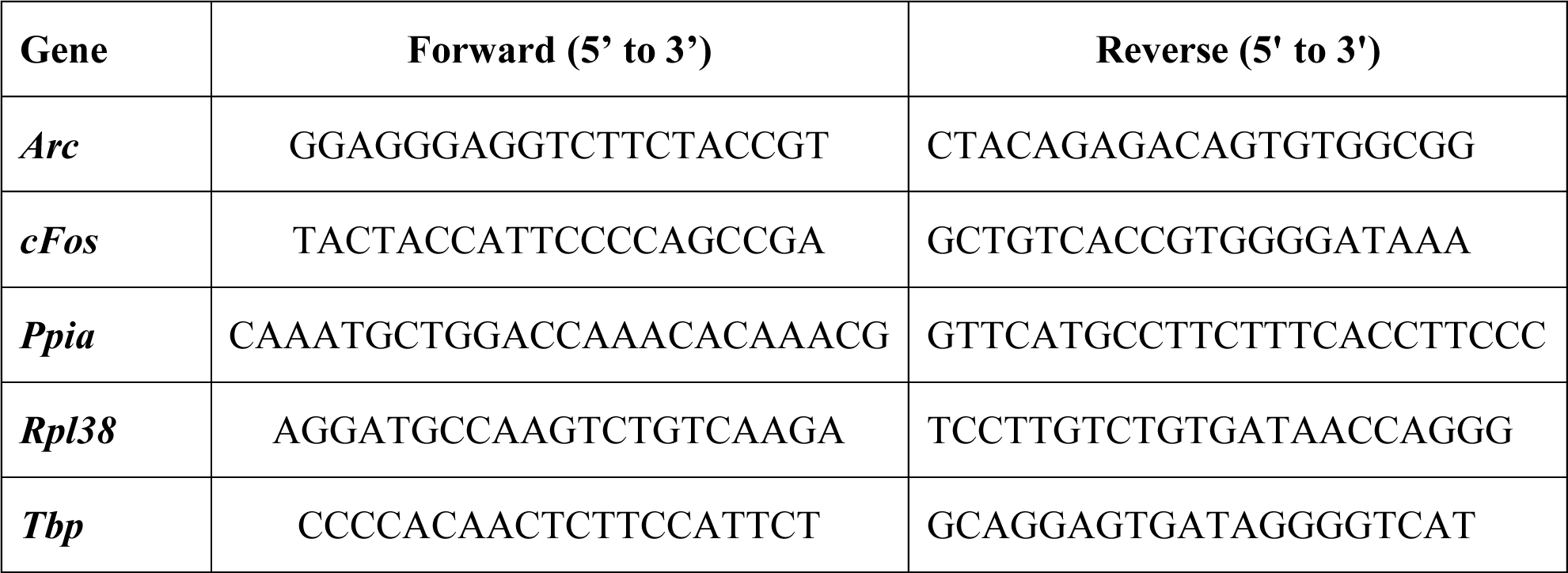
List of primers used for quantitative real-time PCR.

**Table S2-S5 available as separate Excel files:**

**Table S2:** Lists of clusters gene markers.

**Table S3:** Lists of DEGs for GFP-positive vs GFP-negative nuclei.

**Table S4**: Lists of DEGs for *Arc*-positive vs *Arc*-negative nuclei.

**Table S5**: Lists of DEGs for reactivated vs non-reactivated nuclei.

